# Audiovisual processing is selectively impaired in the primary visual cortex but not in the posterior parietal cortex of a mouse model of Fragile X syndrome

**DOI:** 10.1101/2025.04.04.647162

**Authors:** Luca Montelisciani, Conrado A. Bosman, Umberto Olcese

## Abstract

Sensory processing disruptions are a common feature in various forms of autism spectrum disorders (ASD), yet the circuit-level mechanisms underlying these deficits remain poorly understood. Here, we examined differences in audiovisual processing between wild-type and Fmr1-knockout (KO) mice, a model of Fragile X syndrome, which itself represents the most common hereditary cause of autism. We performed multi-area Neuropixels probe recordings in the primary visual cortex (V1) and posterior parietal cortex (PPC) of head-fixed, anesthetized mice being shown visual and auditory stimuli. Previous studies generally reported sensory hypersensitivity in Fragile X mice. In contrast, we observed reduced visual and auditory responses in V1 (but not PPC) of Fragile X compared to wild-type mice. Auditory stimuli reduced the amplitude of visual responses in both V1 and PPC across the two lines, but a dependency on the relative timing of the two sensory modalities was only observed in wild-type mice. Our results provide evidence for a region-specific sensory-processing phenotype of ASD, something that needs to be taken into account when evaluating the efficacy of pharmacological and behavioral strategies to address ASD symptoms.

## Introduction

Sensory processing disruptions are a common feature in various forms of autism spectrum disorders (ASD) (Baum et al., 2015; Robertson and Baron-Cohen, 2017). Such sensory issues tend to hinder nearly all aspects of daily life, with sensory-related symptoms often preceding and predicting later social and behavioral difficulties linked to autism (Baum et al., 2015; Robertson and Baron-Cohen, 2017). Gaining insight into the neural circuit alterations that interfere with typical sensory processing in ASD has therefore been proposed as a key step towards clarifying the mechanisms underlying a wide range of autistic traits and opening avenues for therapeutic possibilities (Baum et al., 2015; El Idrissi and McCloskey, 2023). Fragile X syndrome (FXS) is particularly valuable for studying these mechanisms as it represents the most common hereditary cause of ASD (Freund and Reiss, 1991; Rifé et al., 2003), has been long studied in human subjects (Bagni and Zukin, 2019; Fung and Reiss, 2016), and has a well-established animal model – the Fmr1-KO mouse – that mimics several characteristics of the human condition (Bernardet and Crusio, 2006; Comery et al., 1997; Mineur et al., 2006; Spencer et al., 2005). Previous studies in Fmr1-KO mice have attempted to uncover the circuit-level mechanisms underlying impaired sensory processing. Compared to wild-type (WT) mice, neuronal activity in Fmr1-KO mice is characterized by an imbalanced excitation-inhibition ratio (Antoine et al., 2019; Contractor et al., 2015; Gibson et al., 2008), which results in the hyper-excitability and often hyper-responsiveness of pyramidal neurons located in primary sensory cortices (Domanski et al., 2019a; Gonçalves et al., 2013). Such hyperexcitability is thought to stem from reduced activity in fast-spiking interneurons (Domanski et al., 2019a; Gibson et al., 2008; Goel et al., 2018; Kourdougli et al., 2023). Several studies, however, also reported no difference in either spontaneous or sensory-evoked activity in sensory cortices of Fmr1-KO mice, and instead suggested that hyperexcitability might affect other properties of cortical activity such network synchrony or stimulus selectivity (Arbab et al., 2018; Cheyne et al., 2019; Goel et al., 2018; Kissinger et al., 2020). Thus, the mechanisms underlying sensory processing deficits in FXS are still not fully understood.

Importantly, most studies have so far focused on either spontaneous or sensory-evoked activity in primary sensory cortices only, and were limited to the analysis of unisensory processing. However, primary sensory cortices are only the first step in the cortical hierarchy for processing sensory information. Since many of the cognitive deficits observed in ASD may stem from an impaired ability of cortical areas to transfer and jointly process sensory information (Baum et al., 2015), it is important to go beyond primary sensory cortices. In particular, the array of higher-order areas surrounding the primary visual cortex (V1) in mice has emerged as a prime candidate for studying the circuit-level mechanisms of sensory processing in areas homolog to the ventral and dorsal stream in primates (Andermann et al., 2011; Glickfeld et al., 2014; Khastkhodaei et al., 2016; Marshel et al., 2011; Oude Lohuis et al., 2022a; Wang et al., 2012; Wang and Burkhalter, 2007). Higher-order visual areas have been implicated in several forms of visual processing, complementing and expanding the role of V1 (Huh et al., 2018; Jin and Glickfeld, 2020; Keller et al., 2020; Khastkhodaei et al., 2016; Murgas et al., 2020; Oude Lohuis et al., 2022a; Pak et al., 2020). Furthermore, a key sensory deficit in ASD is related to how multiple sensory modalities are jointly processed (Baum et al., 2015; Foss-Feig et al., 2010; Stevenson et al., 2014). In rodents, cortical multisensory processing has typically been studied at the level of the posterior parietal cortex (PPC), which in mice overlaps with dorsal higher-order visual cortices A and AM. In PPC, multisensory interactions have generally been observed to be integrative, with the different sensory modalities evoking excitatory responses, interacting between them following the classical principles of multisensory integration as described for the superior colliculus (Meijer et al., 2019; Olcese et al., 2013; Stein and Stanford, 2008). Multisensory processing has also been described to occur at the level of the primary visual cortex, but the underlying mechanisms still need to be fully understood. For what pertains auditory-evoked activity in V1, both excitatory as well as inhibitory influences have been observed (Ibrahim et al., 2016; Iurilli et al., 2012; Meijer et al., 2019). However, recent studies have revealed that a large fraction of auditory-evoked activity is actually a form of corollary discharge reaching V1 as a consequence of auditory-evoked, stereotyped, and involuntary motor responses (Bimbard et al., 2023; Oude Lohuis et al., 2024a; Williams et al., 2023).

Surprisingly, despite an ever-growing body of knowledge about (multi)sensory processing across primary and higher-order visual cortices, much less is known about what happens in ASD, specifically in the Fmr1-KO model. Addressing this gap is crucial in order to better characterize the neuron-level correlates of impaired sensory processing in ASD and eventually devise interventional strategies to ameliorate such deficits (Dölen et al., 2007; Goel et al., 2018; Kourdougli et al., 2023). To address this gap, we studied if and how sensory processing is impaired in Fmr1-KO mice compared to WT mice by recording the activity of neurons in V1 and PPC during the presentation of visual, auditory, and audiovisual stimuli. We observed reduced visually-evoked responses in V1 of Fmr1-KO compared to WT mice, alongside a decrease in orientation selectivity. Moreover, when visual stimuli were paired with a sound, visually-evoked responses were reduced in both WT and Fmr1-KO mice; however, only in WT mice was such reduction dependent on the relative timing between auditory and visual stimuli. Finally, while PPC neurons show effects similar to V1 for what pertains to the auditory modulation of visual responses, we did not observe any reduction in visually-evoked responses. Overall, these results enhance our understanding of the neural basis of sensory and multisensory deficits in FXS, offering insights into potential therapeutic approaches for ASD and related neurodevelopmental disorders.

## Results

### 1. Reduced visually-evoked responses in V1 of Fmr1-KO compared to WT mice

To investigate visual processing differences between wild-type (WT) and Fmr1-KO mice, we recorded neural activity from the primary visual cortex (V1) using Neuropixels probes (Jun et al., 2017) in head-fixed, urethane-anesthetized mice (Figure 1A). Anesthesia was used to prevent motor confounds during later auditory stimulation (Bimbard et al., 2023; Oude Lohuis et al., 2024a).

**Figure 1:**
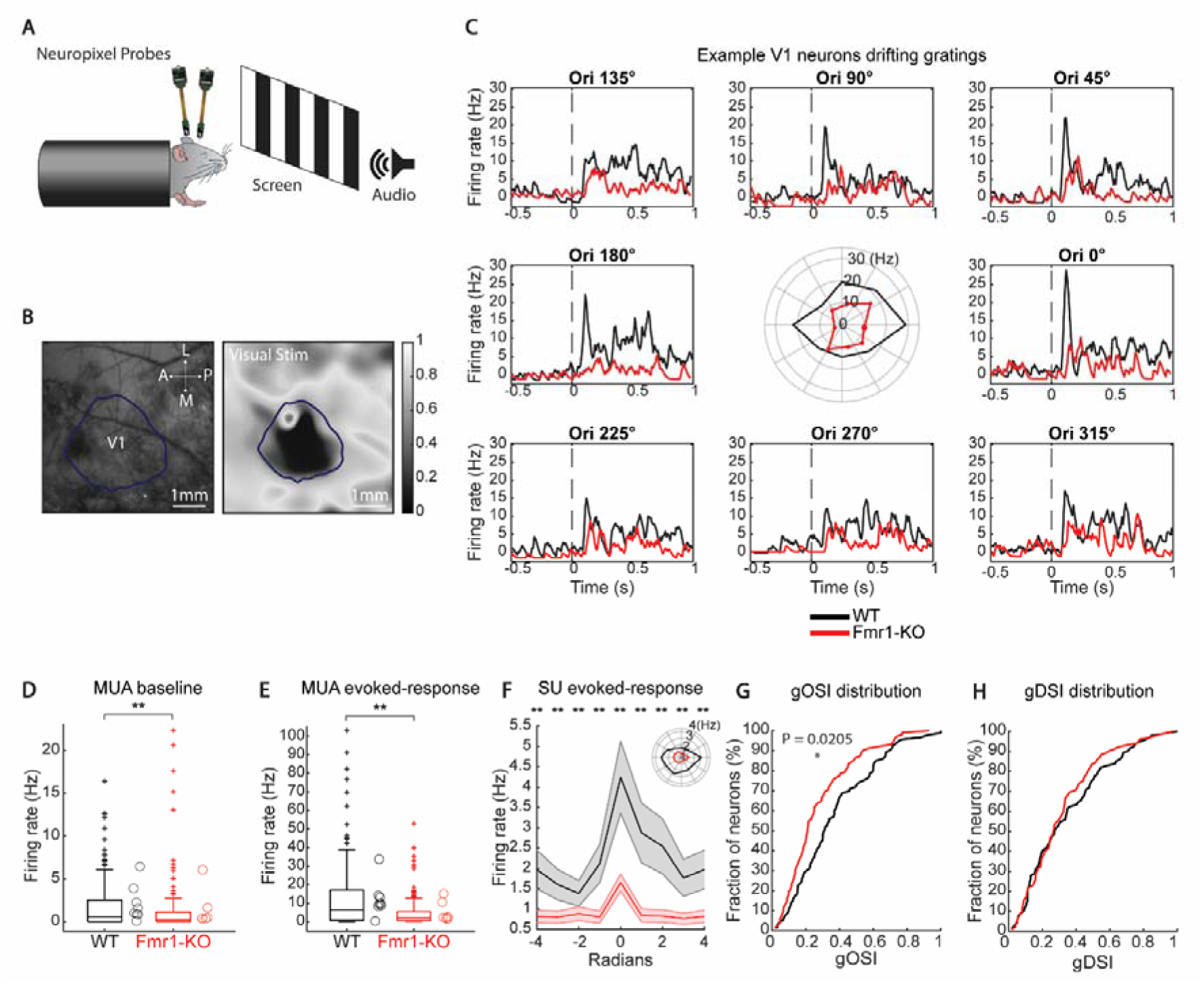
Visually evoked responses in V1 of Fmr1-KO and WT mice. **(A)** Experimental setup. Head-fixed mice were implanted with two Neuropixels probes (one targeting V1, the other PPC), and exposed to visual and auditory stimuli. **(B)** Intrinsic optical signal imaging (IOS). Cortical areas were localized using IOS. The visually evoked signal (right panel) was overlaid on the vessel map (left panel) to identify V1. **(C)** Peri-Stimulus Time Histograms (PSTHs) for two example V1 neurons, one from a WT (black) and one from a Fmr1-KO (red) mouse, during drifting grating presentation. The central polar plot shows the neurons’ tuning curves. **(D)** Mean baseline firing rates for all visually responsive MUA. Each circle represents an individual animal’s average. **(E)** Same as (*D)*, but for responses to the preferred orientation. **(F)** Average tuning curves of V1 neurons for WT and Fmr1-KO mice, aligned to the preferred direction (0 rad). Shaded areas indicate mean ± SEM. Inset: Tuning curve shown in polar coordinates. Asterisks indicate significant differences at specific orientations (*p < 0.05)**. (G)** Cumulative distributions of gOSI for WT and Fmr1-KO (p < 0.005, Kolmogorov–Smirnov test). **(H)** Same as (*G)* for gDSI. (**p < 0.01, *p < 0.05)

We localized V1 using intrinsic optical signal imaging (IOS, Fig 1B; see also Materials and Methods) and confirmed the placement of Neuropixels probes via histological analysis. Visual responses were evoked using drifting gratings along eight different directions (Figure 1C). This allowed the quantification of orientation selectivity and direction selectivity by computing the global orientation (gOSI) and direction selectivity indexes (gDSI), respectively (see also Materials and Methods) (Niell and Stryker, 2008; Oude Lohuis et al., 2022a; Zhao et al., 2013).

During drifting grating presentation, both baseline firing rates and baseline-corrected responses to the preferred orientation of visual-only responsive units were reduced in Fmr1-KO mice compared to WT (p=0.004, Linear mixed-effects model; WT = 202 units, Fmr1-KO = 145 units; Figure 1D-E). This reduction was evident in both multi-unit activity (MUA) and single-unit activity (SU) (Suppl. Figure 1A-B). Given that reduced visually evoked responses in Fmr1-KO mice could be influenced by lower baseline firing rates, we computed z-scored firing rates to account for this effect (see Materials and Methods). Similarly to non-normalized responses, z-scored visually evoked responses were also lower in Fmr1-KO (Suppl. Figure 1C), confirming reductions in both baseline firing rates and sensory-evoked activity.

To further characterize neuronal responses, we analyzed tuning curves for visually responsive single units (WT = 127 units, Fmr1-KO = 115 units). Sensory evoked responses were significantly reduced in Fmr1-KO mice for both preferred and non-preferred orientations compared to WT (p=0.003, Linear mixed-effects model; Figure 1F, see also Suppl. Figure 1D). We then assessed orientation selectivity by computing gOSI and gDSI for each unit. gOSI values were significantly lower in Fmr1-KO mice, (p=0.02, Kolmogorov– Smirnov test, Figure 1G), whereas DSI remained unchanged between groups (p=0.931, Kolmogorov– Smirnov test, Figure 1H).

Overall, V1 neurons in Fmr1-KO mice exhibited reduced firing rates, evoked responses, and orientation selectivity compared to WT.

### 2. Visually-evoked responses are reduced across all cortical layers in Fmr1-KO compared to WT mice

Previous studies reported alterations in specific neuronal subpopulations and disrupted synaptic inputs across cortical layers in Fmr1-KO mice, potentially affecting feedforward sensory processing and orientation selectivity. To determine whether the reduction in response amplitude and selectivity in V1 varies across cortical layers, we estimated recording depth based on multi-unit activity (MUA) power profiles during visual stimulation (Senzai et al., 2019; see Materials and Methods, Figure 2A,B) and categorized neurons into supragranular, granular, and infragranular layers.

**Figure 2:**
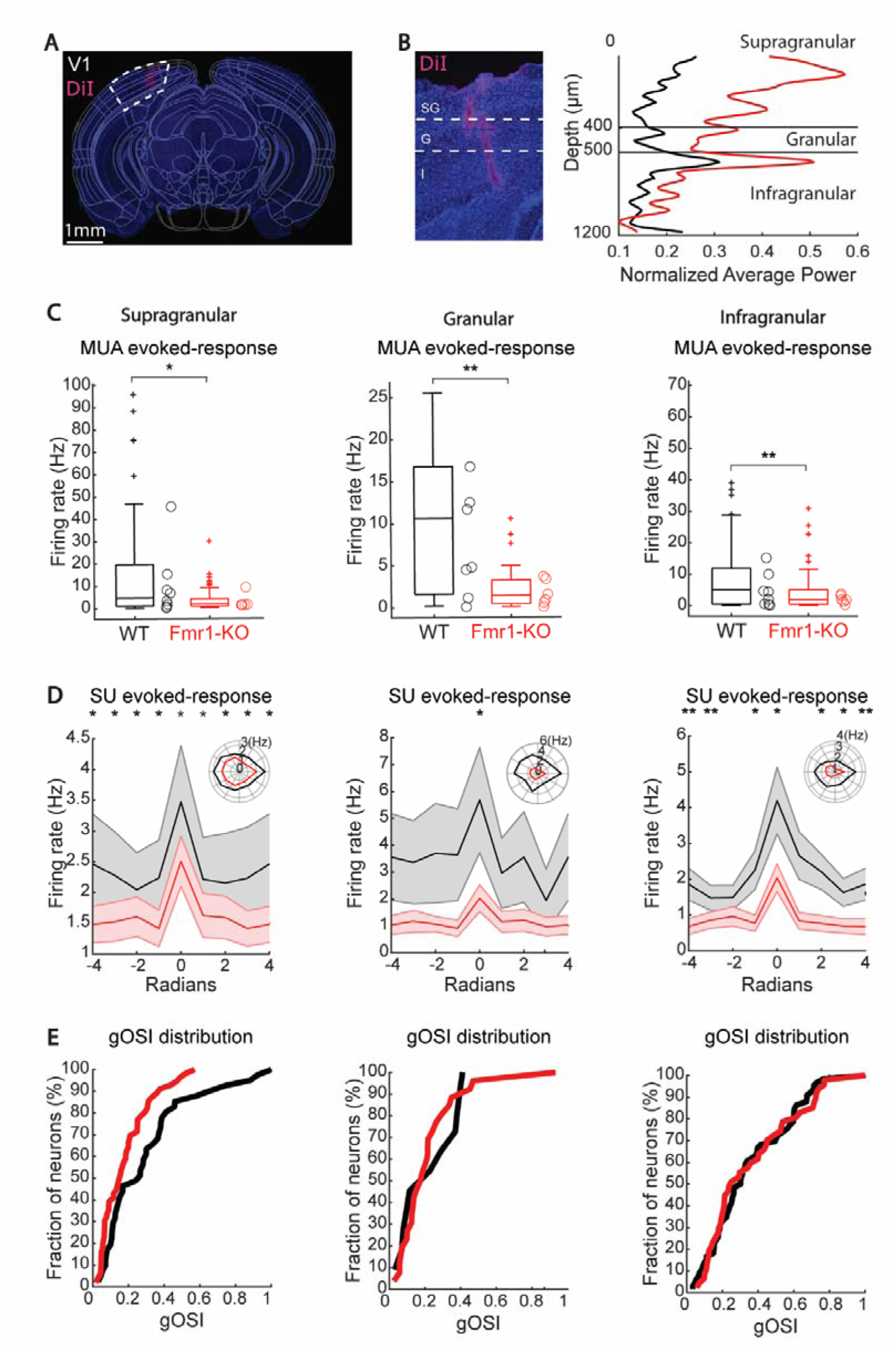
Layer-specific visually evoked responses in V1 of WT and Fmr1-KO mice. **(A-C)** MUA responses to visual stimulation across cortical layers (A: supragranular; B: granular; C: infragranular layers). Circles denote average values per animals. **(D)** Average tuning curves for WT (black) and Fmr1-KO (red) mice across layers. Shaded areas indicate mean ± SEM. Inset: Average response in polar coordinates. **(E)** Cumulative distributions of gOSI for WT (black) and Fmr1-KO (red) mice. (*p < 0.05, ** p < 0.01).

We found that MUA responses to the preferred orientation were significantly lower in Fmr1-KO mice across all layers (supragranular: p=0.038; granular: p=0.004; infragranular: p=0.004; Linear mixed-effects models; WT/Fmr1-KO units per layer: supragranular 70/69, granular 24/29, infragranular 117/75; Figure 2C). We then analyzed visually-evoked activity in SU across both preferred and non-preferred orientations. Fmr1-KO mice exhibited a significant reduction in the amplitude of evoked responses compared to WT mice across all layers (Figure 2D), with the strongest deficits in supragranular and infragranular layers. In contrast, we found a significant reduction in granular neurons only for responses to the preferred orientation. When calculating the gOSI, we found no significant differences between lines for any of the cortical layers (Figure 2E).

Besides laminar alterations, earlier studies also highlighted changes in the activity of parvalbumin-positive interneurons as a key feature in the phenotype of Fmr1-KO mice (Antoine et al., 2019; Contractor et al., 2015; Gibson et al., 2008; Goel et al., 2018; Kourdougli et al., 2023). We thus categorized neurons into putative fast-spiking interneurons and pyramidal neurons based on action potential waveforms (Niell and Stryker, 2008; Olcese et al., 2013; Oude Lohuis et al., 2022a; see Materials and Methods). Surprisingly, we found a larger-than-expected fraction of narrow-spiking neurons (corresponding to putative fast-spiking interneurons; WT/Fmr1-KO: regular-spiking 71/83; fast-spiking 130/61; see Suppl. Figure 2A). Thus, we cannot exclude that the narrow-spiking neurons that we identified might also include putative excitatory units. Baseline activity was similar between genotypes for broad-spiking neurons (p-value 0.070, with a trend toward higher firing rates in Fmr1-KO mice), but was significantly higher in WT for narrow-spiking neurons (p=0.003, Linear mixed-effects model, Suppl. Figure 2B). Visually-evoked responses were similarly reduced in Fmr1-KO compared to WT mice for both broad- and narrow-spiking neurons (p < 0.05, Linear mixed-effects model; Suppl. Figure 2C,D).

Overall, these results indicate that the reduction in visually evoked responses in Fmr1-KO mice is widespread across cortical layers and neuronal subpopulations, pointing towards a generalized reduction in visually evoked responses across all V1 layers.

### 3. Reduced auditory-evoked responses in V1 of Fmr1-KO mice

V1 not only processes visual stimuli but also responds to auditory stimuli (Meijer et al., 2019; Oude Lohuis et al., 2024a; Pennartz et al., 2023). Given reports of cross-modal integration deficits in ASD subjects (Baum et al., 2015; Foss-Feig et al., 2010), we assessed auditory responsiveness in V1 by presenting pure tones at two different frequencies (Fig. 3B).

**Figure 3:**
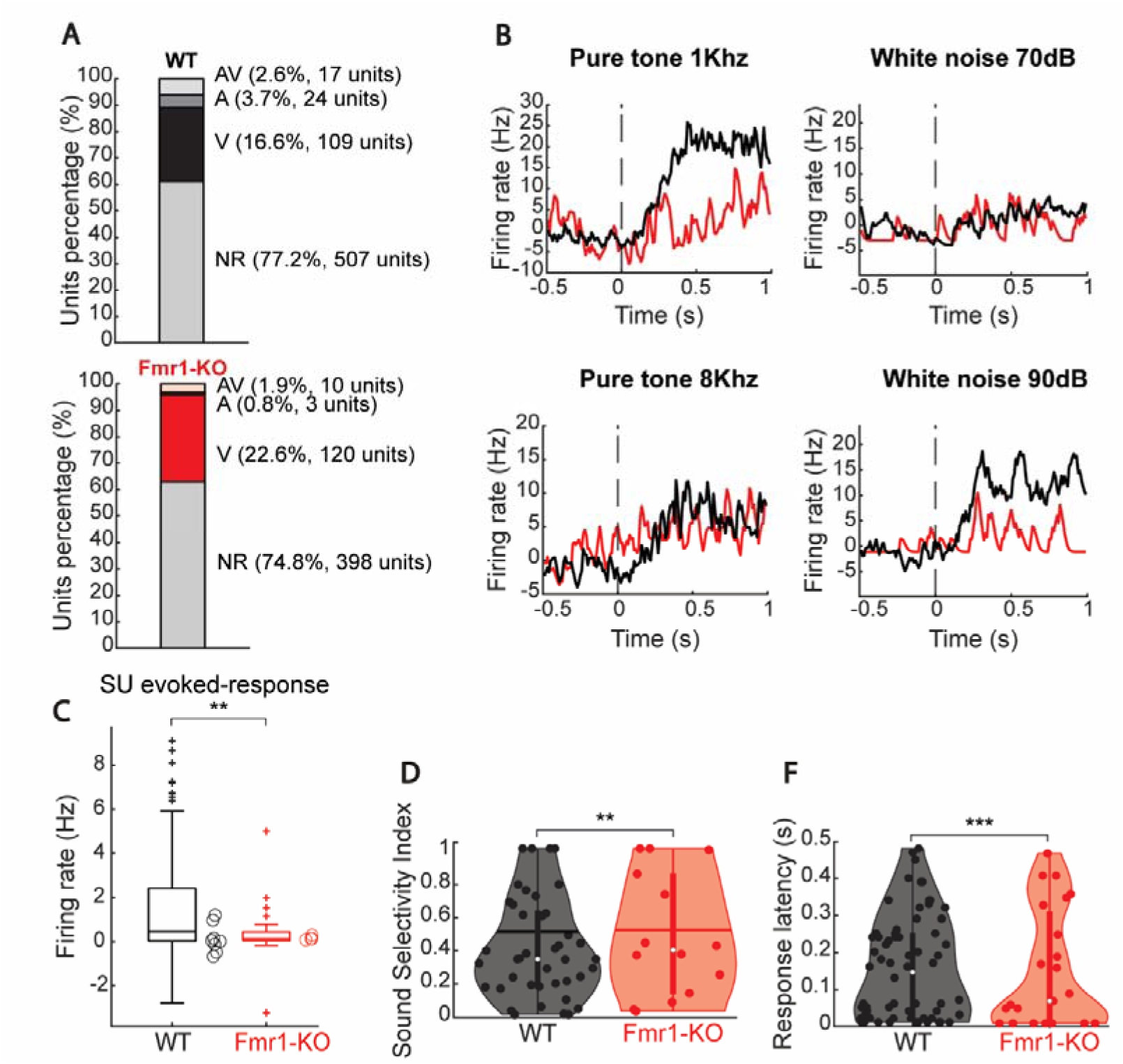
Auditory-evoked activity in mouse V1. **(A)** Fraction of neurons modulated by visual and/or auditory stimuli in WT (top) and Fmr1-KO (bottom) mice. **(B)** Four example V1 single units showing auditory responses to pure tones (1 kHz and 8 kHz) and white noise (70 db and 90 db amplitude) in WT and Fmr1-KO mice (black and red, respectively). **(C)** Amplitude of auditory-evoked V1 responses in SU in WT and Fmr1-KO mice (black and red respectively). Circles denote the average values for single animals. **(D)** Sound selectivity index for auditory-responsive SUs recorded from WT and Fmr1-KO mice (black and red, respectively; each dot represents a SU). **(E)** Same as *D*, but for sound response latency. (* indicates p-value < 0.05, ** indicates p-value < 0.01)

We first determined the fraction of neurons significantly modulated by pure tones. Fewer neurons responded to auditory compared to visual stimuli in both genotypes (WT: 3.7% auditory vs. 16.6% visual, p < 0.0001, Chi-square test; Fmr1-KO: 0.8% auditory vs. 32% visual, p < 0.0001, Chi-square test; Figure 3A). WT mice exhibited a significantly higher fraction of auditory-responsive units compared to Fmr1-KO mice (Chi-square test, p < 0.001). No significant differences were observed when we presented white noise bursts instead of pure tones in a subset of mice (Chi-square test, p = 0.3187; Figure 3B). Auditory evoked responses were markedly reduced in Fmr1-KO compared to WT mice for both MUA and SU, in line with what we observed for visually evoked responses (p = 0.007, Linear mixed-effects model, WT = 114 units, Fmr1-KO = 32 units, Figure 3C).

We further analyzed the sound selectivity index (SSI, see Materials and Methods) to evaluate the specificity of auditory responses to different frequencies. SSI values were significantly higher in WT compared to Fmr1-KO mice (p = 0.003, Linear mixed-effects model, Figure 3D). However, the broad distribution of SSI values in both cohorts suggests that auditory responses in V1 are only moderately selective. Finally, we examined auditory response latencies to distinguish true, short-latency auditory responses from longer-latency auditory-evoked motor correlates (Oude Lohuis et al., 2024a). Response latency was significantly shorter in Fmr1-KO than in WT mice (p = 0.0004, Linear mixed-effects model, Figure 3E), and within the range of both direct sensory responses – i.e., less than about 30 ms in V1 (Oude Lohuis et al., 2024b) – as well as potential residual motor-related activity.

### 4. Auditory stimuli reduce visual response amplitude in both WT and Fmr1-KO mice

Previous studies have shown that, besides directly evoking responses, auditory stimuli can also modulate visual responses in V1 (Ghazanfar and Schroeder, 2006; Ibrahim et al., 2016; Iurilli et al., 2012; McClure and Polack, 2019; Meijer et al., 2017; Williams et al., 2023). To assess whether such modulation differs between WT and Fmr1-KO mice, we examined how auditory stimuli influenced visually evoked responses. Given that the relative timing of sensory inputs is crucial for cross-modal interactions (Lippert et al., 2013; Olcese et al., 2013) and is altered in ASD (Foss-Feig et al., 2010; Stevenson et al., 2014), we paired drifting gratings with pure tones at various stimulus onset asynchronies (SOAs: −250 ms, −150 ms, −50 ms, 0 ms, +50 ms and +150 ms, where negative SOAs indicate that the auditory stimulus was presented first; Figure 4A).

**Figure 4.**
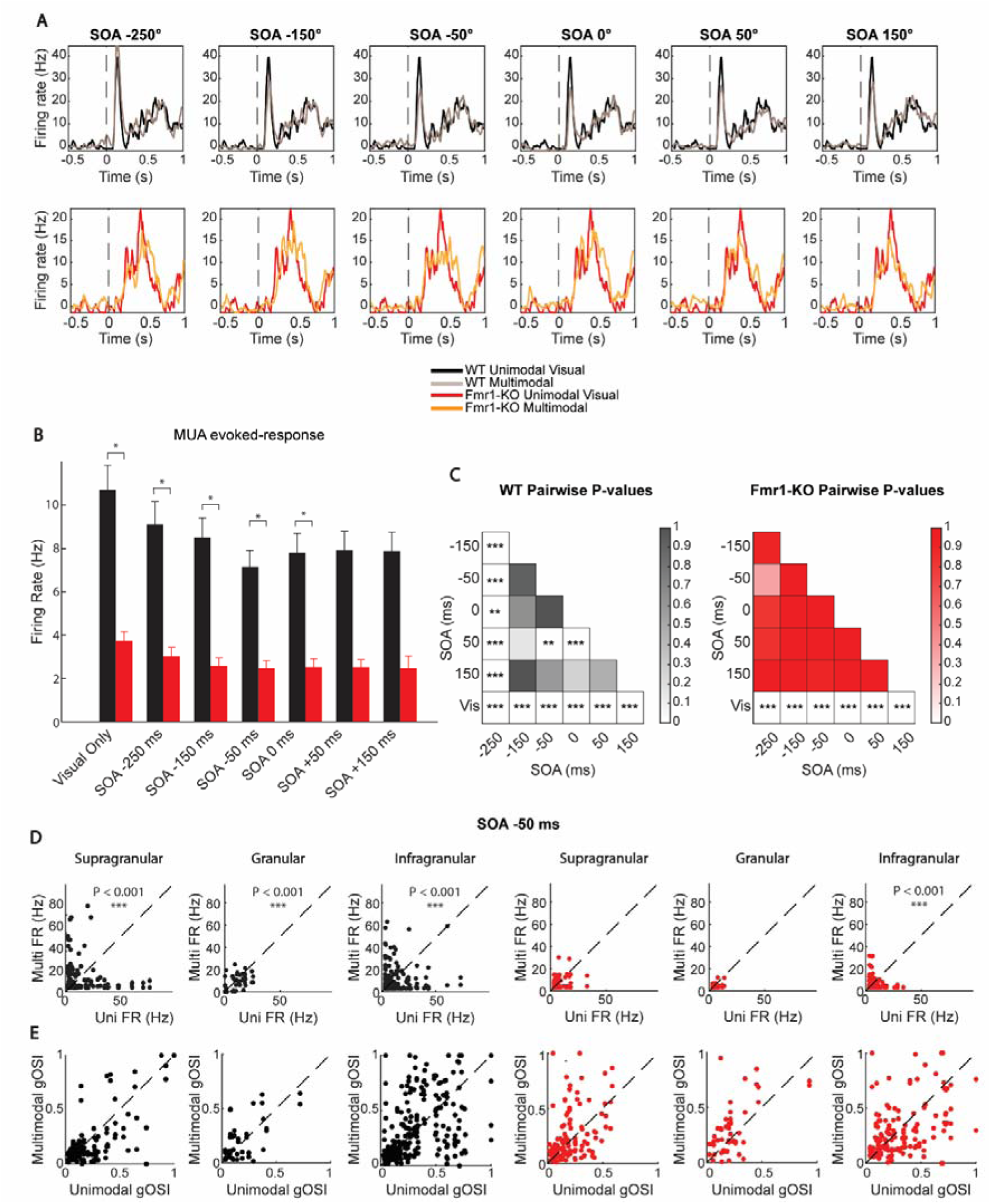
: Auditory modulation of visually-evoked responses in V1 of WT and Fmr1-KO mice. **(A)** Example PSTHs for unimodal and multimodal (i.e., paired with pure tones) visually-evoked responses in WT and Fmr1-KO mice for different SOA. **(B)** MUA visually-evoked responses in WT and Fmr1-KO as a function of SOA. **(C)** Heatmap of significant differences between the conditions depicted in *B*. **(D)** Auditory modulation of SU visual responses across layers for both WT and Fmr1-KO mice. Plots refer to the −50 ms SOA, for which the strongest reduction in response amplitude was observed. See Fig. S3 for all SOA values. **(E)** Same as *D*, but for gOSI. See Fig. S4 for all SOA values. (* indicates p-value < 0.05, ** indicates p-value < 0.01, *** indicates p-value < 0.001)

In both genotypes, visual responses were significantly reduced when paired with auditory stimuli compared to responses to visual stimuli alone (p < 0.05, Two-way mixed-effects ANOVA; WT = 202 units, Fmr1-KO = 145 units; Figure 4B-C). As observed earlier, WT mice generally exhibited higher firing rates than Fmr1-KO mice (Figure 4B) and showed significant SOA-dependent modulation of visual responses, with specific SOAs (e.g., −150 ms and 0 ms) leading to the strongest suppression of visual responses (Figure 4C-D, Suppl. Figure 3). This suppression occurred across all layers in WT mice (Figure 4D, Suppl. Figure 3). In contrast, SOA had no effect on auditory modulation of visual responses in Fmr1-KO mice (Figure 4C). Also, despite reduced visually evoked response amplitude, orientation selectivity remained unaffected by auditory stimuli (Figure 4E, Suppl. Figure 4). Overall, these findings suggest that while sounds suppress visual responses in both genotypes, SOA-dependent modulation is only present in WT mice.

### 5. Limited Fmr1-KO-specific modulation of sensory processing in mouse posterior parietal cortex

V1 is only the first cortical stage of visual processing. Furthermore, its role in auditory processing remains debated (Bimbard et al., 2023; Oude Lohuis et al., 2024b; Pennartz et al., 2023). To investigate the subsequent cortical stages of visual and multisensory processing, we next focused on the posterior parietal cortex (PPC), a key area for integrating sensory information in rodents (Meijer et al., 2019; Olcese et al., 2013; Oude Lohuis et al., 2022b) – Figure 5A. As in V1, we observed both visually and auditory evoked responses (Figure 5B,C). However, PPC had a significantly larger fraction of auditory-responsive neurons compared to V1 (WT: Chi-square test, p < 0.0001; Fmr1-KO: Chi-square test, p < 0.0001, Figure 5I).

**Figure 5:**
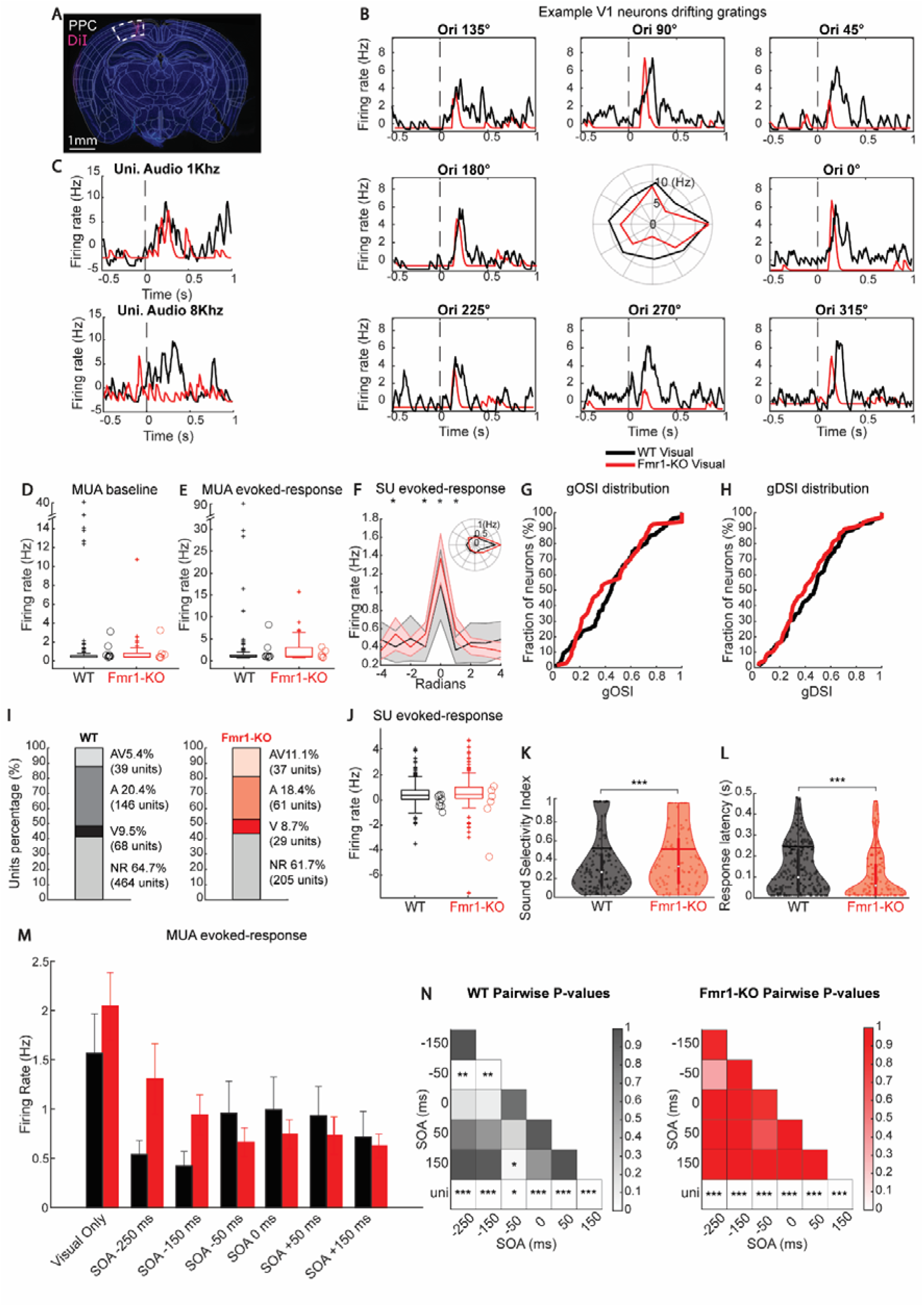
Visual, auditory and audiovisual responses in PPC of WT and Fmr1-KO mice. **(A)** DAPI-stained (blue) coronal section showing DiI-stained (red) electrode track in Posterior parietal cortex (−2.2 mm posterior to bregma). **(B)** PSTHs showing visually-evoked responses to drifting gratings for two example SU in PPC from respectively one WT (black) and one Fmr1-KO (red). The polar plot at the center of the panel shows the tuning curve of the example neurons. **(C)** Same as *B*, but for two example neurons showing responses to auditory stimuli. **(D)** Baseline firing rates for all visually-responsive MUA. Circles denote the average values for single animals. **(E)** Same as *D* but for visually-evoked responses to the preferred orientation for all detected MUA. **(F)** Average tuning curves of PPC SUs for WT and Fmr1-KO mice to gratings drifting along different directions (aligned to the preferred direction, set at 0 rad). Curves with shading indicate mean ± SEM. Inset: Average tuning curve displayed in polar coordinates. Asterisks indicate significant differences between lines for a given direction**. (G,H)** Cumulative distributions for gOSI (left panel) and gDSI (right panel) for WT and Fmr1-KO. **(I)** Fraction of neurons modulated by visual and/or auditory stimuli in WT (top) and Fmr1-KO (bottom) mice. **(J)** Amplitude of auditory-evoked PPC responses in SU in WT and Fmr1-KO mice (black and red respectively). Circles denote the average values for single animals. **(K)** Sound selectivity index for auditory-responsive SUs recorded from WT and Fmr1-KO mice (black and red, respectively; each dot represents a SU). **(L)** Same as *I*, but for sound response latency. **(M)** MUA visually-evoked response in WT and Fmr1-KO as a function of SOA. **(N)** Heatmap of significant differences between conditions depicted in *M*. (* indicates p-value < 0.05, ** indicates p-value < 0.01, *** indicates p-value < 0.001)

Unlike V1, PPC MUA responses showed no differences between WT and Fmr1-KO mice in terms of either baseline or visually evoked activity (Figure 5D,E). SU tuning curves revealed some significant differences (Figure 5F), with visually evoked responses higher in Fmr1-KO PPC neurons compared to WT. However, no significant differences were observed in either gOSI or gDSI (Figure 5G,H). Auditory evoked responses also did not differ between genotypes (Figure 5J). Interestingly, in Fmr1-KO mice, auditory responses were more selective to specific sound frequencies compared to WT mice, as reflected by a higher sound selectivity index (p=0.0001; Linear mixed-effects model; Figure 5K). Additionally, PPC neurons in Fmr1-KO mice exhibited significantly longer auditory response latencies compared to WT (p=0.0001, Linear mixed-effects model, Figure 5L).

Finally, we assessed whether auditory stimuli modulated visual responses in PPC. As in V1, sounds reduced visual response amplitudes ad the MUA level (p=0.0001, Linear mixed-effects model, Figure 5M,N, see Suppl. Figure 5 for SU results; number of MUA/SU: WT 129/107, Fmr1-KO 96/66). Similarly to V1, SOA-specific modulations were only observed in WT mice (Figure 5M,N), although only between specific SOA values. Overall, in contrast to V1, PPC showed limited differences between WT and Fmr1-KO mice for what pertains visual, auditory, and audiovisual responses.

## Discussion

Our findings provide a comprehensive characterization of visual, auditory and multisensory processing deficits in Fmr1-KO mice. In V1 (but not PPC), we observed a generalized reduction in the amplitude of visual and auditory-evoked responses, which were accompanied by a reduced feature selectivity, in particular for what pertains the orientation of visual stimuli. The amplitude of visual responses was reduced when auditory stimuli were simultaneously presented. In both V1 and PPC of WT (but not Frm1-KO) mice, this effect depended on the relative timing of auditory and visual stimuli, although this effect was more prominent in V1.

The reduction in the amplitude and orientation selectivity of visually evoked responses that we observed in V1 underlies a possible microcircuit-level basis for the deficits in sensory processing that have been identified as a key hallmark of ASD (Baum et al., 2015; Robertson and Baron-Cohen, 2017). In particular, weaker and more broadly tuned neuronal responses might underlie perceptual deficits such as difficulty in determining the directionality of noisy visual stimuli (Robertson et al., 2014; Spencer et al., 2000). Analogously to visual responses, auditory-evoked responses in V1 were also dampened in Fmr1-KO compared to WT mice. Importantly, similar effects were observed across V1 layers (Fig. 2). Thus, the reduction is response amplitude and selectivity appears to be a generalized phenotype in V1 of Fmr1-KO mice. Several previous studies identified sensory hypersensitivity as a key trait in the somatosensory cortex of Fmr1-KO mice (Domanski et al., 2019a; Gibson et al., 2008; He et al., 2017; Kourdougli et al., 2023). However, this might be a trait specific to the somatosensory system of Fmr1-KO mice that does not necessarily extend to other sensory modalities. In fact, no change in visually-evoked activity was previously observed in V1 of Fmr1-KO compared to WT mice (Goel et al., 2018; Kissinger et al., 2020). The reduction in the amplitude of sensory-evoked responses in V1 might be explained by several factors. First, we performed recordings under urethane anesthesia to prevent auditory-related motor responses (Bimbard et al., 2023; Oude Lohuis et al., 2024b); this might have affected sensory responses and their dependency of the overall excitation/inhibition balance (Albrecht and Davidowa, 1989; Sceniak and MacIver, 2006; Shumkova et al., 2021). Second, repeated presentation of visual stimuli has been shown to induce a stronger reduction in the amplitude of oscillatory responses in Fmr1-KO compared to WT mice (Kissinger et al., 2020); this might have been reflected in our study by a reduction in spiking responses already within a single recording session. Thus, the results that we obtained are not necessarily in contrast with previous findings, but might rather expand our understanding of how sensory processing differs in FXS. More broadly, previous studies evidenced a reduction in interneuronal activity, in particular for fast-spiking parvalbumin-positive interneurons, as a key hallmark of the FXS phenotype in Fmr1-KO mice (Domanski et al., 2019a; Gibson et al., 2008; Goel et al., 2018). In line with this, we also observed a reduction in baseline firing rates in narrow-spiking, but not broad-spiking interneurons (Suppl. Figure 2B).

This reduction in inhibitory tone has been classically linked to the sensory hypersensitivity reported in FXS; however, a reduction in the activity of parvalbumin-positive interneurons may also paradoxically impair the reliability and response fidelity of excitatory neurons (Kato et al., 2017; Zhu et al., 2015) and is therefore also compatible with a reduction in trial-averaged evoked responses. Overall, our result thus showcase how sensory processing is disrupted in FXS, starting from basic features such as amplitude and feature selectivity of evoked responses (Goel et al., 2018).

Besides unisensory processing, we also explored how multisensory processing differed in Fmr1-KO compared to WT mice. Deficits in multisensory processing, in fact, are thought to be a hallmark of the ASD phenotype (Baum et al., 2015), but the underlying neuron-level mechanisms are poorly understood. In line with previous studies (Iurilli et al., 2012), we observed that auditory stimuli reduced the amplitude of visual responses (Figure 4). Such effect was observed across both mouse lines, but only in WT mice was it dependent on SOA. This aligns with evidence suggesting that the temporal window for multisensory integration is broadened in ASD (Foss-Feig et al., 2010; Stevenson et al., 2014). Thus, our results support the view that deficits in multisensory processing, and specifically in the ability to bind multisensory features over time (Brock et al., 2002; Stevenson et al., 2014), are a key feature of the FXS phenotype – and by extension ASD (Baum et al., 2015).

Surprisingly, in contrast to V1, we found similar correlates of unisensory and multisensory processing between WT and Fmr1-KO mice in PPC. This suggests that sensory processing might be differentially affected across cortical areas in Fmr1-KO mice. It is also possible that deficits in (multi)sensory processing might be limited to primary sensory areas – although to our knowledge ours is the first study to investigate sensory processing correlates in Fmr1-KO mice outside such areas. Nevertheless, mouse PPC has been hypothesized to be a cortical hub for multisensory processing, receiving information from multiple sensory areas and playing a crucial role in the multisensory decision making process (Bizley et al., 2016; Meijer et al., 2019; Olcese et al., 2013; Wang et al., 2012). Thus, in view of the predominance of (multi)sensory processing deficits in ASD, it is surprising that we observed limited differences in PPC responsiveness between the two mouse lines. In view of the large cortical network for multisensory processing in mice (Meijer et al., 2019), however, it might be that other association cortices show instead Fm1-KO-specificic deficits in sensory processing. Another factor to consider is that PPC itself shows neural correlates of audiovisual processing but no causal involvement in the detection of such stimuli (Oude Lohuis et al., 2022b); therefore, its role in determining sensory processing deficits in ASD might analogously also be limited.

Our study suffered from some limitations. First, as mentioned earlier, we were limited to probing only two cortical areas (V1 and PPC), something that prevented us from uncovering the full scope of a potential area-specific sensory processing phenotype. Furthermore, recordings were done under anesthesia, a condition that is known to affect the transfer of information between cortical areas (Ishizawa et al., 2016; Mashour, 2024) – even if short-range projections such as those between V1 and PPC have been shown to be mostly preserved across wakefulness and anesthesia (Olcese et al., 2013; Oude Lohuis et al., 2022a). We cannot therefore exclude that sensory and multisensory processing will be differentially affected in other cortical areas and in the awake state. Related to this, the absence of a behavioral paradigm also prevented us from linking specific changes in sensory processing to any behavioral consequence, and thus directly connect neuronal activity to sensory perception. For what pertains auditory-evoked responses in V1, these have long been described in mouse V1 (Iurilli et al., 2012). However, recent studies suggested that a large fraction of such auditory-evoked activity actually stems from auditory-induced motor responses (Bimbard et al., 2023; Oude Lohuis et al., 2024b) eliciting widespread cortical activity (Stringer et al., 2019). In order to remove such confounding factors, we therefore performed our experiments under anesthesia. Still, a sizable fraction of auditory-evoked activity showed latencies more compatible with a motor rather than a sensory origin – cf. Figure 3, (Oude Lohuis et al., 2024b). Thus, we cannot exclude that a fraction of the auditory-evoked activity we observed might only indirectly have an auditory origin. However, our results still indicate that all the forms of evoked activity that we observed in V1 show a reduced amplitude in Fmr1-KO compared to WT mice. Finally, our study was only correlative, and did not explore the mechanistic link between specific neuronal subpopulations (or area) and multisensory processing (Goel et al., 2018). Addressing this is imperative in order to determine which circuit-level manipulations can restore sensory processing and ultimately perception in Fmr1-KO mice, paving the way for therapeutic interventions.

In summary, our study demonstrates that Fmr1-KO mice exhibit distinct unisensory and multisensory processing deficits, with impairments ranging from basic visual and auditory responses to multisensory integration dynamics. Such deficits are pervasive in V1, but surprisingly not in the closely connected association area PPC. Overall, these findings not only provide a circuit-level framework for better understanding sensory challenges in FXS, but also offer potential avenues for developing therapies aimed at restoring normal sensory network function. Addressing these sensory processing deficits may have far-reaching implications for improving quality of life in individuals with FXS, ASD and related neurodevelopmental disorders.

## Acknowledgments

This work was supported by the FLAG-ERA JTC 2019 project DOMINO (co-financed by NWO) and the FLAG-ERA JTC 2023 project MONAD (co-financed by NWO) to C.A.B. and U.O. The authors would like to thank Gerjan Huis in ‘t Veld for his support to experimental activities.

## Materials and Methods

### Subjects

All animal experiments were performed according to national and institutional regulations. The experimental protocol was approved by the Dutch Commission for Animal Experiments and by the Animal Welfare Body of the University of Amsterdam (Ethical protocol number 2020-10466). A total of 14 mice (both female and male) were used: 8 WT (C57BL/6j) and 6 Fmr1-KO mice. WT mice included 3 male and 5 female animals. Fmr1-KO mice were bred on a homozygous background (KO/KO) and crossed with SST-Cre and VIP-Cre transgenic lines to generate Fmr1-KO;SST-Cre and Fmr1-KO;VIP-Cre offspring. Crossing were originally planned for an experiment involving optogenetic modulation of interneuronal activity, which is not the subject of the current study. The final cohort consisted of 3 Fmr1-KO;SST-Cre (1 male, 2 female) and 3 Fmr1-KO;VIP-Cre (1 male, 2 female) mice. The Fmr1-KO line was originally developed by The Dutch-Belgian Fragile X Consortium (1994, Cell). Animals were at least five weeks of age at the start of experiments. Mice were group-housed, with ad libitum access to water and food, under a normal day–night schedule.

### Experimental Design

#### P0 Viral Injections

Neonatal Cre mice (P0, Fmr1-KO;SST-Cre and Fmr1-KO;VIP-Cre only) were anesthetized by hypothermia, placed on a chilled surface, and maintained under ice anesthesia throughout the procedure. AAV viral vectors (AAV2.1 – DIO – GyFP – ChR2) were injected unilaterally (left hemisphere) into the target brain region using a glass micropipette attached to a Nanoject pressure injection system (Drummond Scientific Company). The injection was performed at a controlled volume to ensure proper viral diffusion while minimizing tissue damage. Following the injection, neonates were placed under heating lamps until they regained normal movement and responsiveness before being returned to their home cage.

#### Headbar Implantation

Initially, mice were administered urethane (20% solution, 0.8 ml/hg I.P.) and allowed to rest for at least 30 minutes. Anesthesia depth was confirmed by the absence of reflexive responses to paw pinch. The skull was then shaved, disinfected, and a circular area excised, with the remaining skin were secured using tissue adhesive (3-M Vetbond, 3M, St. Paul, MN, USA). A custom-fabricated titanium head-bar with a 10 mm circular recording chamber was positioned over the exposed left hemisphere, covering visual, auditory, and somatosensory cortices. The head-bar was affixed with cyanoacrylate and C&B Super-Bond (Sun Medical), to ensure stability for neurophysiological recordings.

#### Intrinsic Optical Imaging

Before conducting intrinsic optical imaging, we thinned the skull to enhance the optical access to the visual cortex. To localize the primary visual cortex, we performed intrinsic optical imaging (IOS) under urethane anesthetized conditions (Olcese et al., 2013; Oude Lohuis et al., 2022b). A vasculature image was acquired under 540 nm light before starting the imaging session. During IOS, the cortex was illuminated with monochromatic 630 nm light. Images were acquired at 1 Hz using the Labeotech’s OiS200 Modular Optical Imaging System (250 × 250 pixels) defocused ∼500–600 μm below the pial surface. We presented visual stimuli, consisted of full-field drifting gratings (spatial frequency 0.05 cpd, temporal frequency 1.5 Hz) for 1 s in each of eight directions. We acquired 6 s of baseline signal and 12 s of hemodynamic response during stimulation. The acquired frames during the response were baseline-subtracted, averaged, and thresholded to produce a map of V1. PPC was localized based on the location of higher order visual areas A and AM with respect to V1 (Oude Lohuis et al., 2022b),and was situated approximately 1.5–2.0 mm anterior and 1.5–2.0 mm medial relative to the center of V1.

#### Craniotomies

After IOS, we performed a small craniotomy based on the image acquired using a dental drill (200 μm) over the areas of interest (V1 and PPC). The dura was then removed in order to eliminate any possible impediment for probe insertion.

#### In Vivo Electrophysiology

Mice were fixated in a custom-built holder in a dark and sound-attenuated cabinet. The headbar was attached to a custom-made holder via two screws. Recordings were performed under urethane. Body temperature was kept constant through a heating pad at 37.5°C. Extracellular recordings were performed with Neuropixels 1.0 probe (Jun et al., 2017). In each recording session, the electrode arrays were slowly inserted until the recording sites spanned the cortical layers. Prior to insertion, electrodes were dipped in DiI (ThermoFisher Scientific) allowing better post-hoc visualization of the electrode tract. After insertion, the exposed cortex and skull were covered with saline. The probe ground was connected to the headbar and the reference electrode to the saline solution. Recordings started at least 15 min after insertion to allow for tissue stabilization.

#### Sensory Stimulation

Visual stimuli were generated in custom-written MATLAB routines (The MathWorks) with Psychtoolbox extensions (Brainard). Visual stimuli were gamma-corrected and presented with a 60-Hz refresh rate on an 18.5-inch monitor positioned at a 45° angle with the body axis from the mouse at 21 cm from the eyes, subtending 91° horizontally and 60° vertically. Visual sensory stimulation consisted of drifting gratings in eight directions (0°–360°, in 45° steps), presented in a pseudo-random order, with a spatial frequency of 0.04 cycles/degree and a temporal frequency of 2 Hz at 100 % contrast.

Auditory stimuli consisted of pure tones (1–81kHz, 11s duration, 701dB intensity) and white noise bursts (1 s burst at 70 dB or 90 db) presented bilaterally from speakers placed at a distance of 20 cm from the mouse.

Auditory and visual stimuli were presented alone or in combination in a pseudorandomized order, separated by a random inter-trial interval from 1.5-2 s. Each condition was presented 20 times. The combined audiovisual stimuli were presented at various SOAs in which the stimuli were presented either at −250, −150, −50, 0, 50, 150 ms apart. Negative values indicate that the auditory stimulus was presented before the visual one.

#### Histology

At the end of the experiment, mice were overdosed with pentobarbital and perfused with 4% paraformaldehyde in phosphate-buffered saline, and their brains were recovered for histology. We cut coronal 50-μm sections with a vibratome, stained them with DAPI (0.3 μM), and imaged mounted sections to verify the location of the recording sites.

### Data Analysis

#### Spike Sorting

Spike detection and sorting were done using Kilosort 3 (Pachitariu et al., 2023). Manual curation was done via the Phy GUI (Rossant et al., 2016). During manual curation, each putative single unit was inspected based on its waveform, autocorrelation function, and firing pattern across channels and time. Only high-quality single units (SU) were included that satisfied the following constraints: 1) less than 0.1% of their spikes within the refractory period of 1.5 ms, and 2) spiking activity was present in at least 20% of the trials. Multiunit activity (MUA) was defined as the set of remaining clusters that did not meet the criteria for SU classification. Specifically, clusters with noisy waveforms, a lack of clear refractory period, or spatially diffuse spike patterns were excluded from the SU pool and classified as MUA—unless they clearly reflected artifacts or non-neuronal signals (e.g., periodic noise, movement-related artifacts), in which case they were discarded entirely. This classification was done by visual inspection during manual curation in Phy, and included clusters that exhibited heterogeneous or overlapping waveforms and consistent firing patterns across trials

#### Classification of Neuron Subtypes

Putative pyramidal and putative fast-spiking interneurons were separated based on the peak-to-trough delay of their average normalized action potential waveform (Niell and Stryker, 2008; Oude Lohuis et al., 2022c). The peak-to-trough delay was computed as the time between peak positive and peak negative voltage deflection (in ms), and single units with a delay lower than 0.50 ms were classified as narrow-spiking, while units with a delay higher than 0.50 ms were classified as broad-spiking. The 0.5 ms cutoff was determined through visual inspection of the distribution of peak-to-trough delays for units identified in WT mice, where a clear separation between the two groups could be observed (Suppl. Figure 2A).

#### Laminar Depth Estimation

The laminar depth of each electrode was estimated based on the depth profile of MUA power (Senzai et al., 2019). Multiunit activity was computed as follows: (i) raw signals were band-pass filtered between 500 Hz and 5,000 Hz using a 100th-order FIR filter to isolate the multiunit activity (MUA) frequency band; (ii) for each trial, 1-second windows following stimulus onset were extracted and averaged across trials to obtain the MUA envelope; (iii) power spectra were computed for each channel using Welch’s method, and the average power was normalized across channels; (iv) the channel closest to the peak of normalized MUA power was manually selected and assigned a nominal depth of 600 µm, designating it as the center of cortical layer 5. Depths of the remaining channels were interpolated assuming 20 µm vertical spacing per channel pair and assigned to supragranular, granular and infragranular layers based on their position relative to the center of layer 5.

#### Sensory-evoked responses

To compute firing rates in response to visual and auditory stimuli, spike times were binned to create Peri-Stimulus Time Histograms (PSTHs) and convolved with a Gaussian window (50-ms standard deviation). Z-scoring was performed by subtracting, for each trial, the mean firing rate of the baseline period (−1 to −0.2 s before stimulus onset) and dividing by the standard deviation of all baseline periods. Only neurons showing a significant sensory-evoked response were retained for further analyses. For visual stimuli, neurons were classified as visual units if their average firing rate in the 0 to 0.5 s post-stimulus window exceeded the mean plus two times the standard deviation of the baseline firing rate (−0.9 to −0.30s) for at least one orientation. Similarly, for auditory stimuli, neurons were classified as auditory units if they exhibited an average response in the 0 to 0.5 s post-stimulus window greater than the mean plus two times the standard deviation of the baseline firing rate (−0.9 to −0.30s) for at least one pure tone.

#### Quantification of Tuning Curves

Tuning curves for single neurons were quantified after computing, independently for each direction the firing rate response to a visual stimulus. The preferred orientation/direction was determined as the one eliciting the largest average firing rate. To align tuning curves, the preferred orientation of each neuron was set to 0 rad and other orientations were displayed relative to this.

#### Orientation and Direction Selectivity

Orientation and direction selectivity were computed using a global orientation selectivity index (gOSI) and a global direction selectivity index (gDSI) (Ibrahim et al., 2016; Ringach et al., 2002). These two measures were computed as follows:

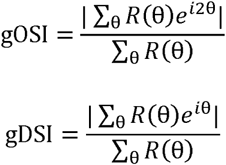

Here, R(θ) is the baseline-corrected firing rate response of a neuron to a bar moving along direction θ and *i* is the imaginary unit. gOSI and gDSI vary between 0 and 1, with 0 indicating a neuron completely untuned for orientation/direction and 1 a neuron only responding to a single orientation/direction, respectively. For the analysis of gOSI and gDSI, only neurons that exhibited significant sensory-evoked responses to at least one stimulus direction were included. Neurons that did not respond to any stimulus direction, defined as the absence of action potential firing, had undetermined gOSI and gDSI values. Consequently, these neurons were excluded from further analysis.

#### Statistical Analysis

Unless specified otherwise, all statistics were performed using linear mixed models (LMMs) in RStudio (RStudio, Boston, MA). LMMs account for the hierarchical structure of our data (e.g., neurons and trials nested within individual mice) and describe the relationships between a response variable and explanatory variables. These models include two types of terms: fixed effects, which represent variables of interest, and random effects, which account for variability within grouping factors (e.g., mouse ID). For all analyses involving hierarchical data, LMMs were constructed with mouse ID as a random effect (intercept only). In analyses where variability between mice was key (e.g., analyses with cohort as a fixed effect), mouse ID was not included as a random effect. Firing rates, which are typically not normally distributed, were analyzed using LMMs without z-scoring. Statistical significance of fixed effects was determined using F-tests with the Satterthwaite approximation to estimate denominator degrees of freedom. Results with a p-value < 0.05 were considered significant. For multiple comparisons, pairwise contrasts were calculated in R. Post hoc pairwise tests were performed to assess the interaction between factors (e.g., SOA and line) using least-squares means with Tukey’s adjustment. For comparisons between groups involving categorical data, such as the number of units between groups, chi-square tests were used. Details of all the statistical analyses that we performed are presented in Supplementary Table 1.

## Data and Software Availability

Original data and the MATLAB, and R scripts used to perform the analyses presented in this manuscript are available by reasonable request to Umberto Olcese.

## Supplementary Figures

**Supplementary Figure 1:**
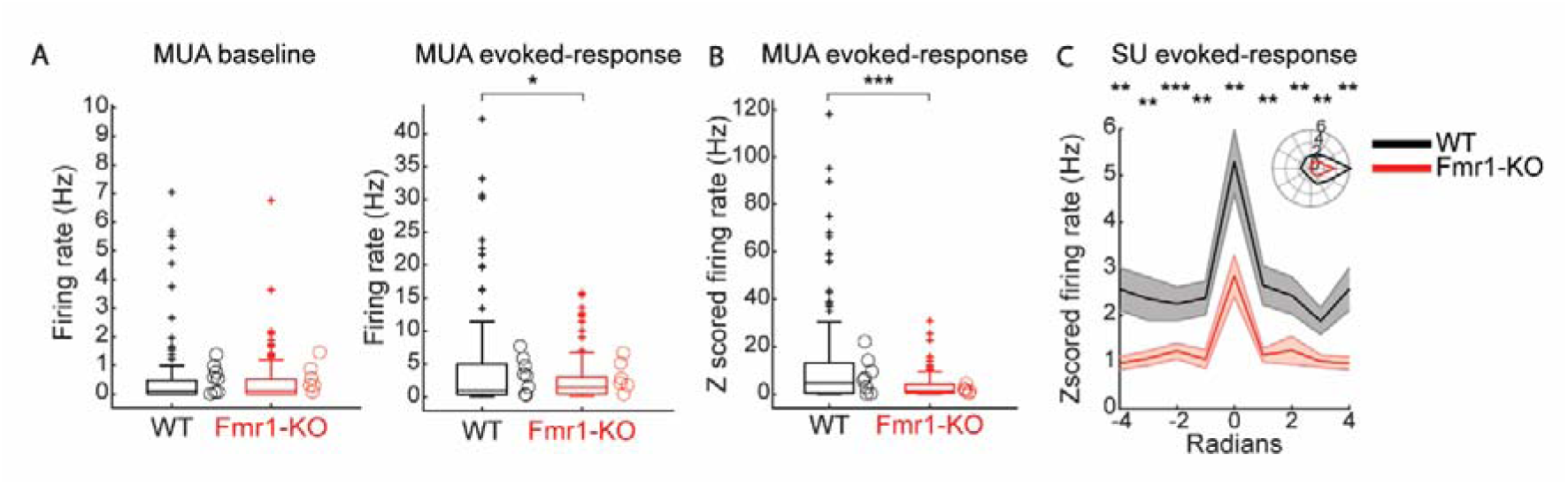
**(A)** Baseline firing rates for all visually-responsive SU. Circles denote the average values for single animals. (**B**) Same as A but for visually-evoked responses to the preferred orientation for all detected SU. (**C**) Same as B but for z-scored values. (**D**) Average tuning curves of V1 neurons z-scored for WT and Fmr1-KO mice to gratings drifting along different directions (aligned to the preferred direction, set at 0 rad). Curves with shading indicate mean ± SEM. Inset: Average tuning curve displayed in polar coordinates. Asterisks indicate significant differences between lines for a given orientation. (* indicates p-value < 0.05, ** indicates p-value < 0.01, *** indicates p-value <0.001)

**Supplementary Figure 2:**
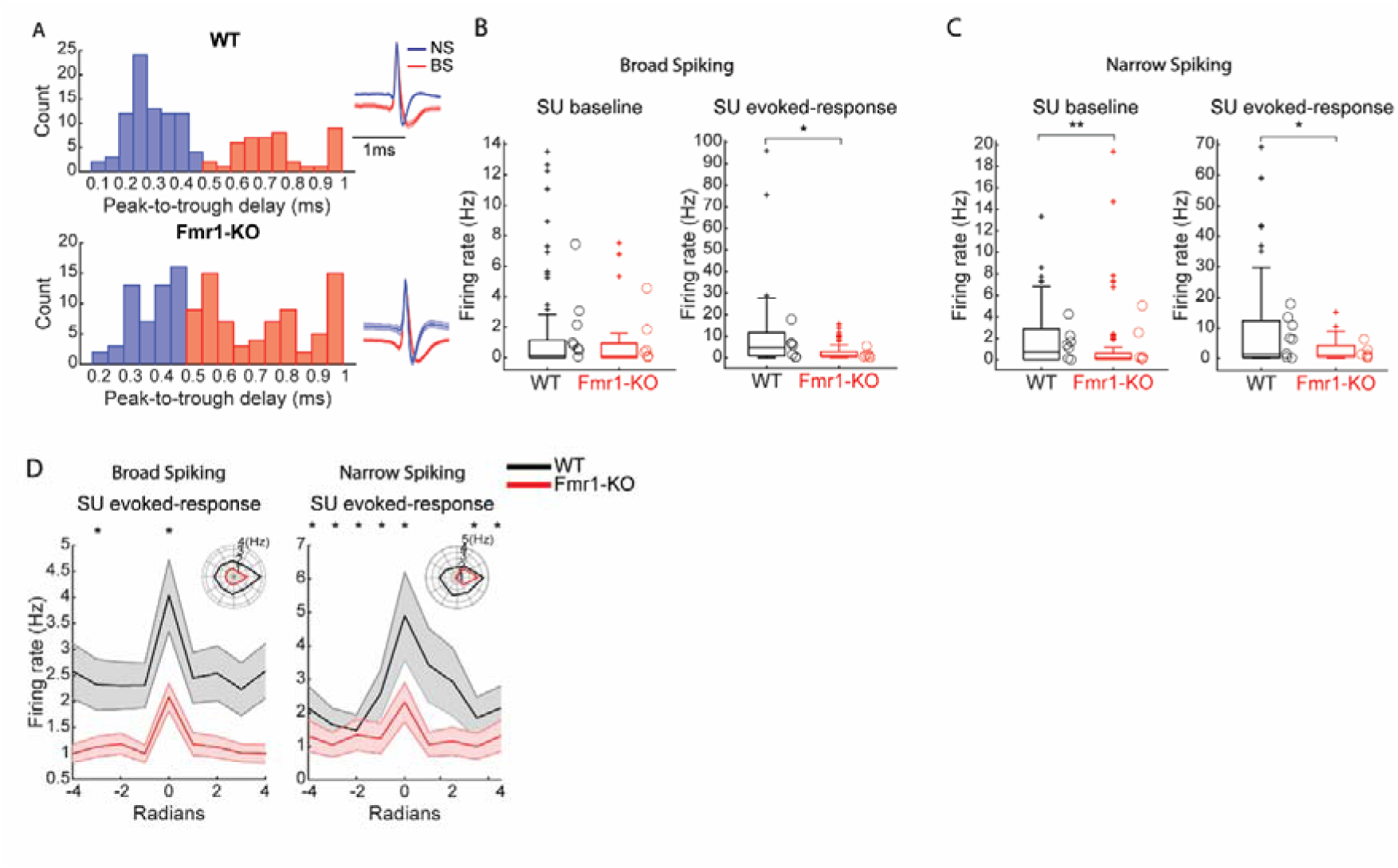
**(A)** Histogram of peak-to-trough delay for all neurons for WT and Fmr1-KO colored by cell type class: narrow-spiking (peak-to-trough delay <0.5 ms; putative inhibitory; blue) and broad-spiking (peak-to-trough delay >0.5 ms; putative excitatory; red). The peak-to-trough delay was capped at 1 ms for neurons whose trough extended beyond the sampled window. **(B**) Baseline and evoked response firing rates for all visually-responsive SU for the broad spiking. Circles denote the average values for single animals. (**C**) Same as B but for narrow spiking. (**D**) Average z-scored tuning curves of V1 neurons for WT and Fmr1-KO mice broad and narrow spiking to gratings drifting along different directions (aligned to the preferred direction, set at 0 rad). Curves with shading indicate mean ± SEM. Inset: Average tuning curve displayed in polar coordinates. Asterisks indicate significant differences between lines for a given orientation. (* indicates p-value < 0.05, ** indicates p-value < 0.01, *** indicates p-value <0.001).

**Supplementary Figure 3:**
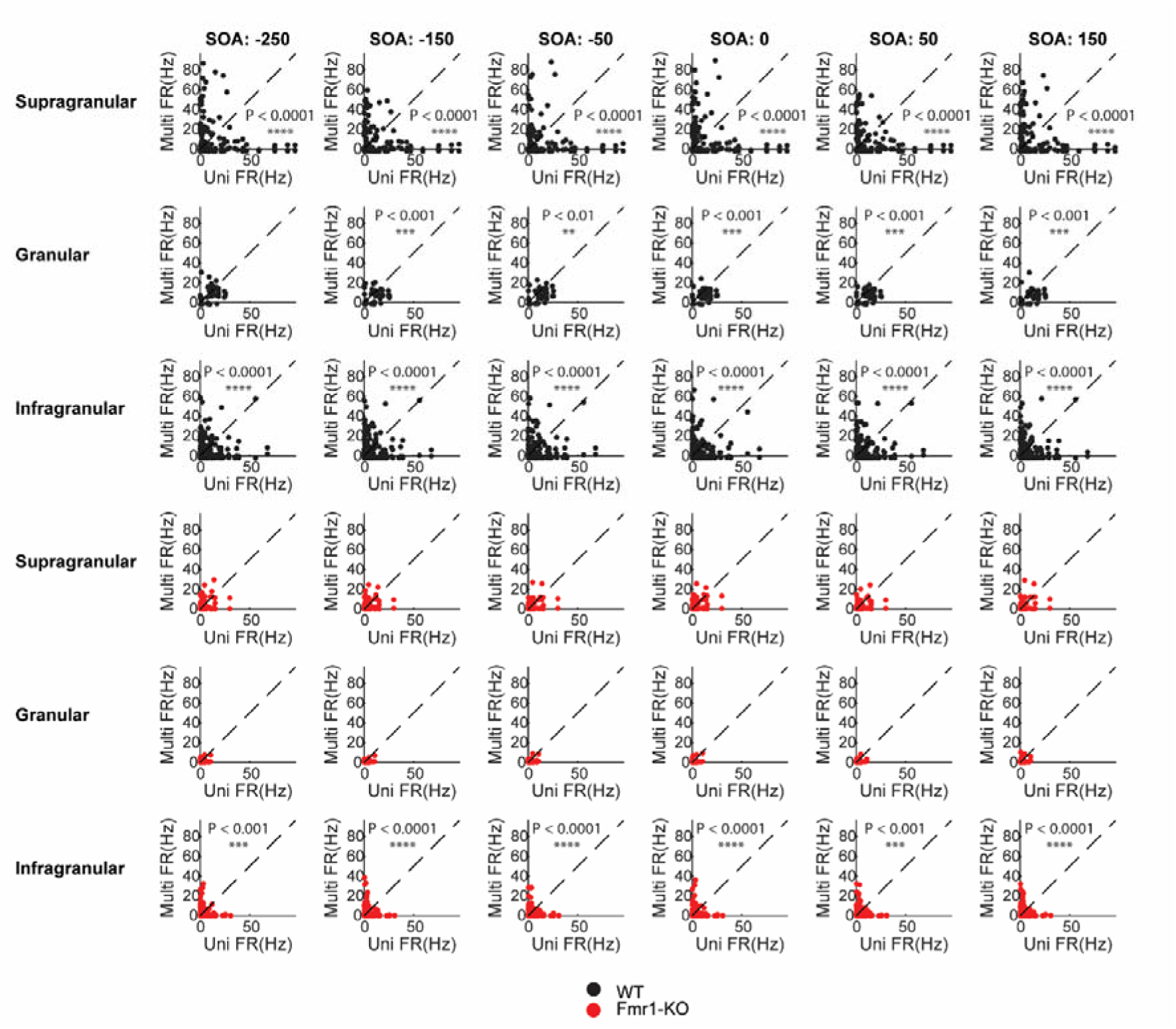
Auditory modulation of V1 SU visual responses across layers for both WT and Fmr1-KO mice and for each SOA (* indicates p-value < 0.05, ** indicates p-value < 0.01, *** indicates p-value < 0.001, **** indicates p-value < 0.00001).

**Supplementary Figure 4:**
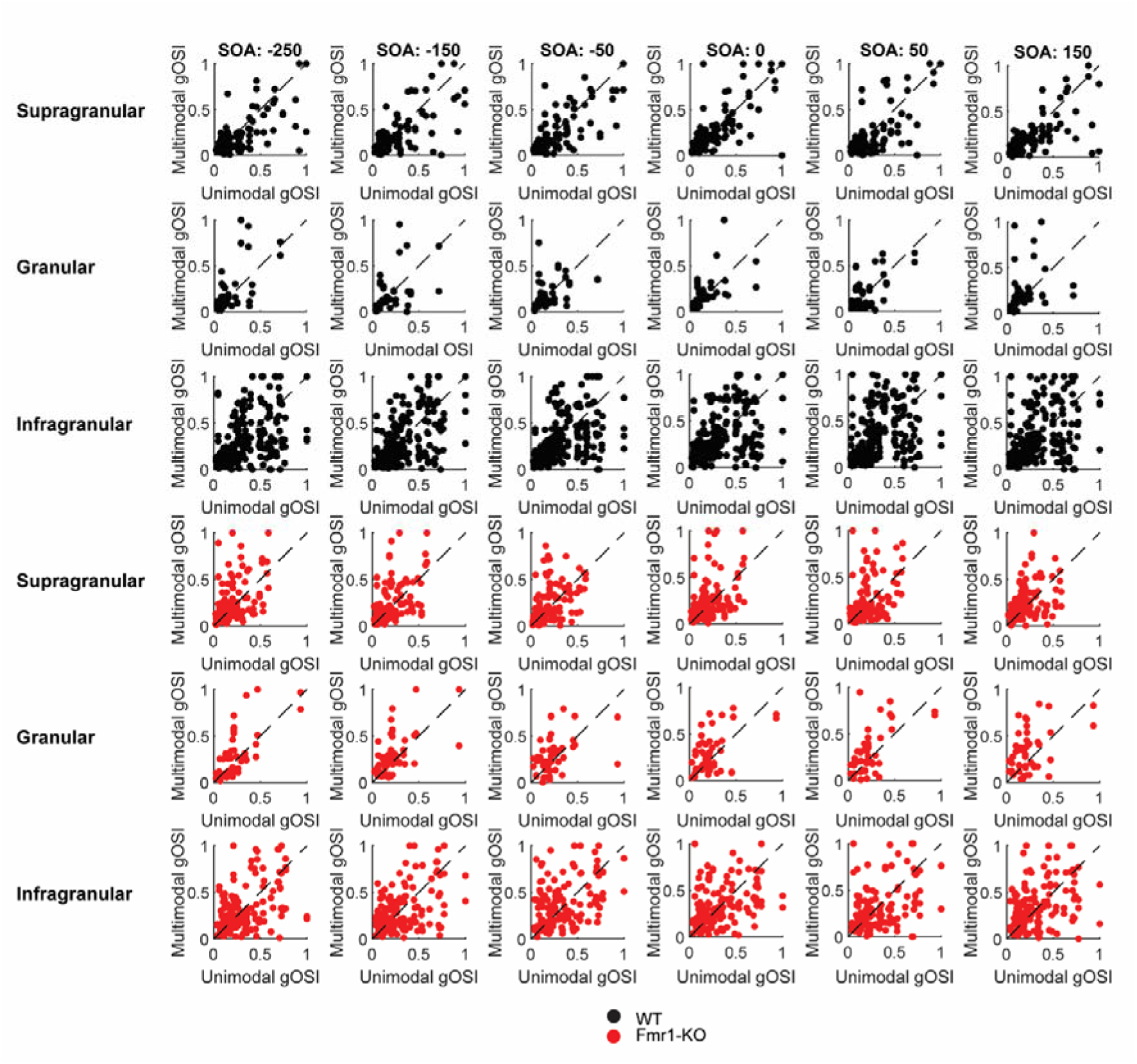
gOSI modulation of V1 SU visual responses across layers for both WT and Fmr1-KO mice and for each SOA (* indicates p-value < 0.05, ** indicates p-value < 0.01, *** indicates p-value < 0.001, **** indicates p-value < 0.00001).

**Supplementary Figure 5:**
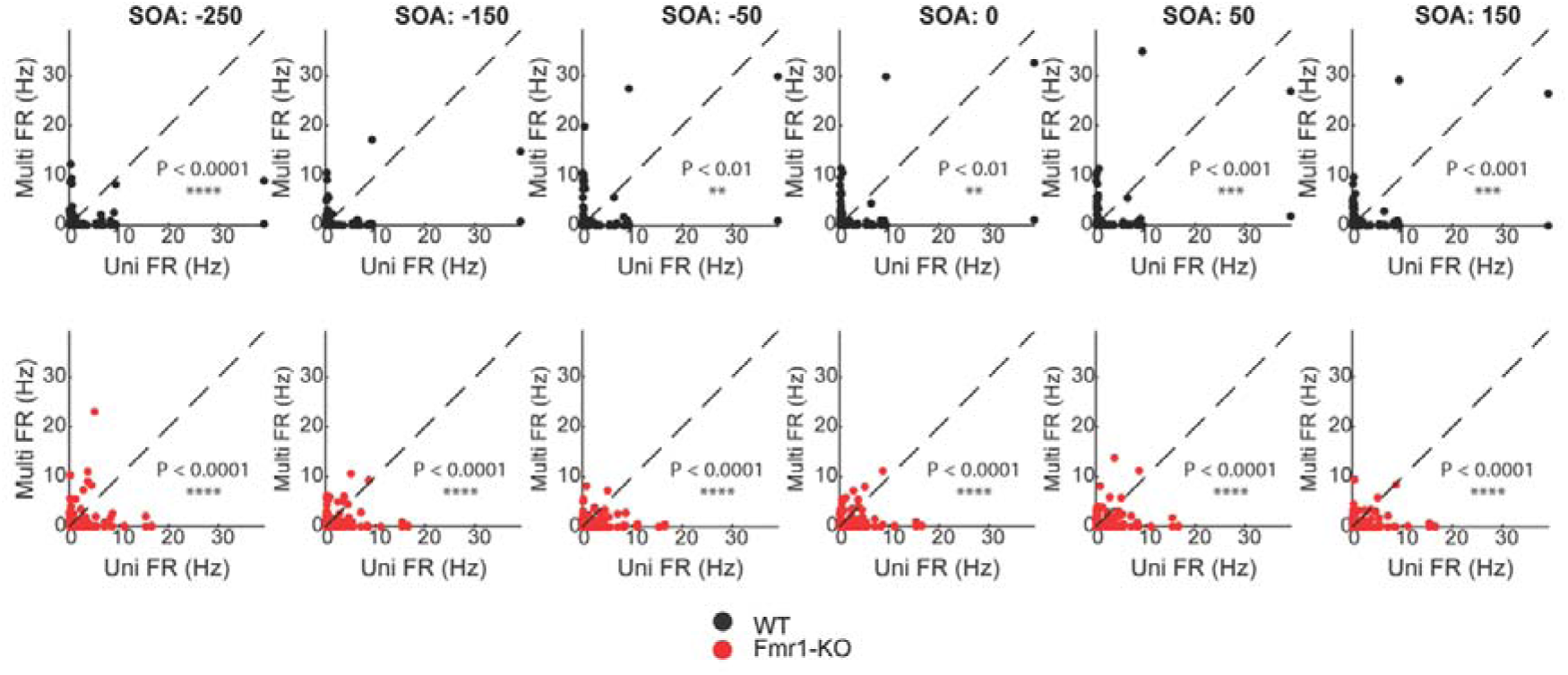
Auditory modulation of PPC SU visual responses across layers for both WT and Fmr1-KO mice and for each SOA (* indicates p-value < 0.05, ** indicates p-value < 0.01, *** indicates p-value < 0.001).

**Supplementary Table 1:** Details of all the performed statistical analyses.

| Figure | Goal | Test | Sample size | Test statistic | P-value |
| --- | --- | --- | --- | --- | --- |
| 1d | Compare baseline firing rates (MUA) | Linear mixed-effects model with Line as a fixed effect and Animal as a random effect | WT = 202 units, Fmr1-KO = 145 units | $F(2, 11.722) = 8.872$ | $p = 0.004$ |
| 1e | Compare visually-evoked responses (MUA) | Linear mixed-effects model with Line as a fixed effect and Animal as a random effect | WT = 202 units, Fmr1-KO = 145 units | | $p = 0.004$ |
| 1f | Compare tuning curves (SU responses) | Linear mixed-effects model with Line as a fixed effect and Animal as a random effect | WT = 202 units, Fmr1-KO = 145 units | $F(2, 8.284) = 10.407$ | $p = 0.003$ |
| 1g | Compare gOSI between groups | Kolmogorov–Smirnov test | WT = 202 units, Fmr1-KO = 145 units | $p = 0.02$ | $p = 0.02$ |
| 1h | Compare gDSI between groups | Kolmogorov–Smirnov test | WT = 202 units, Fmr1-KO = 145 units | $p = 0.931$ | $p = 0.931$ |
| 2c | Compare MUA visually-evoked responses across layers | Linear mixed-effects model with Line as a fixed effect and Animal as a random effect | Supragranular<br>WT = 70 units, Fmr1-KO = 69 units;<br>Granular<br>WT = 24 units, Fmr1-KO = 29 units;<br>Infragranular<br>WT = 117 units, Fmr1-KO = 75 units | S: $F(2, 12.253) = 4.318$<br>G: $F(2, 10.765) = 9.326$<br>I: $F(2, 12.342) = 8.577$ | Supragranular $p = 0.038$; Granular $p = 0.004$; Infragranular $p = 0.004$ |
| 2d | Compare SU evoked responses across orientations and layers | Linear mixed-effects model with Line as a fixed effect and Animal as a random effect | Supragranular<br>WT = 70 units, Fmr1-KO = 69 units;<br>Granular<br>WT = 24, Fmr1-KO = 29;<br>Infragranular<br>WT = 117, Fmr1-KO = 75 | S: $F(2, 10.389) = 6.426$<br>G: $F(2, 8.352) = 4.854$<br>I: $F(2, 7.930) = 5.263$ | Significant for all layers, granular significant only for preferred orientation |
| 2e | Compare gOSI distributions across cortical layers | Kolmogorov–Smirnov test | Supragranular<br>WT = 70 units, Fmr1-KO = 69 units;<br>Granular<br>WT = 24 units, Fmr1-KO = 29 units;<br>Infragranular<br>WT = 117 units, Fmr1-KO = 75 units |  | Not significant for all layers |
| 3a | Compare proportion of auditory-responsive neurons | Chi-square | WT = 657 units, Fmr1-KO = 531 units | | $p\text{-value} = 0.004$ |
| 3c | Compare auditory-evoked SU responses | Linear mixed-effects model with Line as a fixed effect and Animal as a random effect | WT = 114 units, Fmr1-KO = 32 units | $F(2, 12.785) = 7.465$ | $p\text{-value} = 0.007$ |
| 3d | Compare Sound Selectivity Index (SSI) | Linear mixed-effects model with Line as a fixed effect and Animal as a random effect | WT = 114 units, Fmr1-KO = 32 units | $F(2, 7.826) = 12.865$ | $p\text{-value} = 0.003$ |
| 3f | Compare auditory response latency | Linear mixed-effects model with Line as a fixed effect and Animal as a random effect | WT = 114 units, Fmr1-KO = 32 units | $F(2, 8.652) = 22.03$ | $p\text{-value} = 0.004$ |
| 4b | Compare MUA visually-evoked responses with and without auditory modulation across | Two-way mixed-effects ANOVA with Line and SOA as fixed effects, Animal/ID_neuron as random effects | WT = 202 units, Fmr1-KO = 145 units | Line $F(2, 11.5) = 31.749$<br>SOA $F(6, 4452) = 1.750$<br>Line × SOA Interaction | $p < 0.05$ |
| | SOAs | | | $F(6, 4452) = 2.433$ | |
| **4c** | Assess significant differences in visual responses across SOAs in WT and Fmr1-KO | Post hoc pairwise comparisons (lsmeans) | WT = 202 units,<br>Fmr1-KO = 145 units |  | Various (see heatmap) |
| **4d** | Compare SU visual responses with auditory modulation across layers at SOA -50 ms | Two-way mixed-effects ANOVA with Line and Layer as fixed effects, Animal/ID_neuron as random effects | WT = 202 units,<br>Fmr1-KO = 145 units | S: Line $F(2, 11.5) = 31.749$<br>SOA $F(6, 4452) = 1.750$<br>Line × SOA Interaction $F(6, 4452) = 2.433$<br>G: Line $F(2, 11.45) = 17.114$<br>SOA $F(6, 1656) = 0.989$<br>Line × SOA Interaction $F(6, 1656) = 2.226$<br>I: Line $F(2, 11.81) = 43.847$<br>SOA $F(6, 2292) = 1.402$<br>Line × SOA Interaction $F(6, 2292) = 3.172$ | $p < 0.001$ for: WT supra-granular-infra Fmr1-KO for: infra. |
| **4e** | Compare gOSI with and without auditory modulation across layers | Two-way mixed-effects ANOVA with Line and Layer as fixed effects, Animal/ID_neuron as random effects | WT = 202 units,<br>Fmr1-KO = 145 units | S: Line $F(2, 10.93) = 16.132$<br>SOA $F(6, 1140) = 1.495$<br>Line × SOA Interaction $F(6, 1140) = 2.800$<br>G: Line $F(2, 6.43) = 20.224$<br>SOA $F(6, 432) = 3.932$<br>Line × SOA Interaction $F(6, 432) = 1.149$<br>I: Line $F(2, 10.02) = 61.940$<br>SOA $F(6, 1560) = 0.598$<br>Line × SOA Interaction $F(6, 1560) = 1.627$ | Not significant |
| **5d** | Compare baseline firing rates in PPC | Linear mixed-effects model with Line as a fixed effect and Animal as a random effect | WT = 164 units, Fmr1-KO = 73 units | F(2, 2.255) = 2.517 | Not significant |
| **5e** | Compare visually-evoked responses in PPC | Linear mixed-effects model with Line as a fixed effect and Animal as a random effect | WT = 164 units, Fmr1-KO = 73 units | F(2, 4.211) = 2.635 | Not significant |
| **5f** | Compare SU tuning curves for visual responses | Linear mixed-effects model with Line as a fixed effect and Animal as a random effect | WT = 164 units, Fmr1-KO = 73 units | Rad 0: F(2, 7.633) = 6.629 | Significant for rad -3, -1, rad 0, rad 1 (p < 0.05) |
| **5g** | Compare gOSI distributions for visual responses | Kolmogorov–Smirnov test | WT = 164 units, Fmr1-KO = 73 units |  | Not significant |
| **5h** | Compare gDSI distributions for visual responses | Kolmogorov–Smirnov test | WT = 164 units, Fmr1-KO = 73 units |  | Not significant |
| **5i** | Compare n of auditory and visual units in PPC | Chi-square | WT = 717 units, Fmr1-KO = 332 units |  | Not significant |
| **5j** | Compare auditory-evoked responses in PPC | Linear mixed-effects model with Line as a fixed effect and Animal as a random effect | WT = 185 units, Fmr1-KO = 98 units | F(2, 1.242) = 2.540 | Not significant |
| **5k** | Compare sound selectivity index (SSI) in auditory responses | Linear mixed-effects model with Line as a fixed effect and Animal as a random effect | WT = 185 units, Fmr1-KO = 98 units | F(2, 281) = 191.69 | p-value < 0.001 |
| **5l** | Compare auditory response latency in PPC | Linear mixed-effects model with Line as a fixed effect and Animal as a random effect | WT = 185 units, Fmr1-KO = 98 units | F(2, 8.939) = 73.465 | p-value < 0.001 |
| **5m** | Compare MUA visually-evoked responses in PPC across SOAs | Linear mixed-effects model with Line as a fixed effect and Animal as a random effect | WT = 164 units, Fmr1-KO = 73 units | Not significant | Not significant |
| **5n** | Assess significant differences in visual responses across SOAs | Post hoc pairwise comparisons (lsmeans) | WT = 164 units, Fmr1-KO = 73 units |  | WT unimodal vs SOA all significant (p-value < 0.0001) with other significant SOA combination<br>Fmr1-KO unimodal vs SOA all significant (p-value < 0.0001) |

