## Supplementary figures and images for "Audiovisual processing is selectively impaired in the primary visual cortex but not in the posterior parietal cortex of a mouse model of Fragile X syndrome"

### Supplemental Figure 1

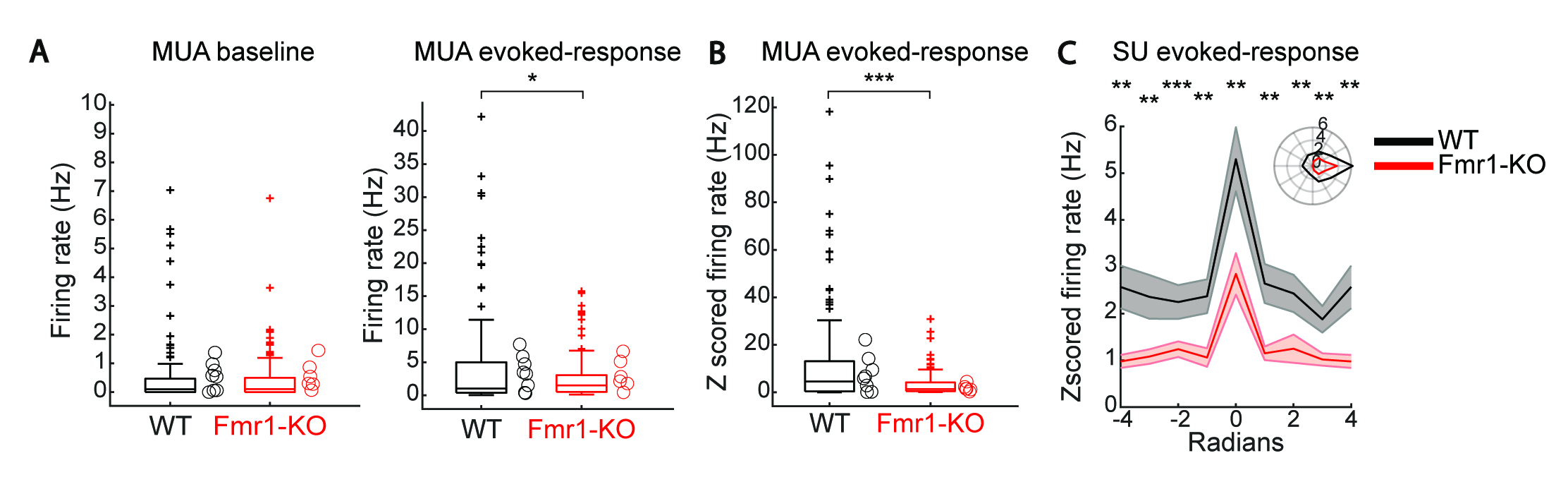

### Supplemental Figure 2

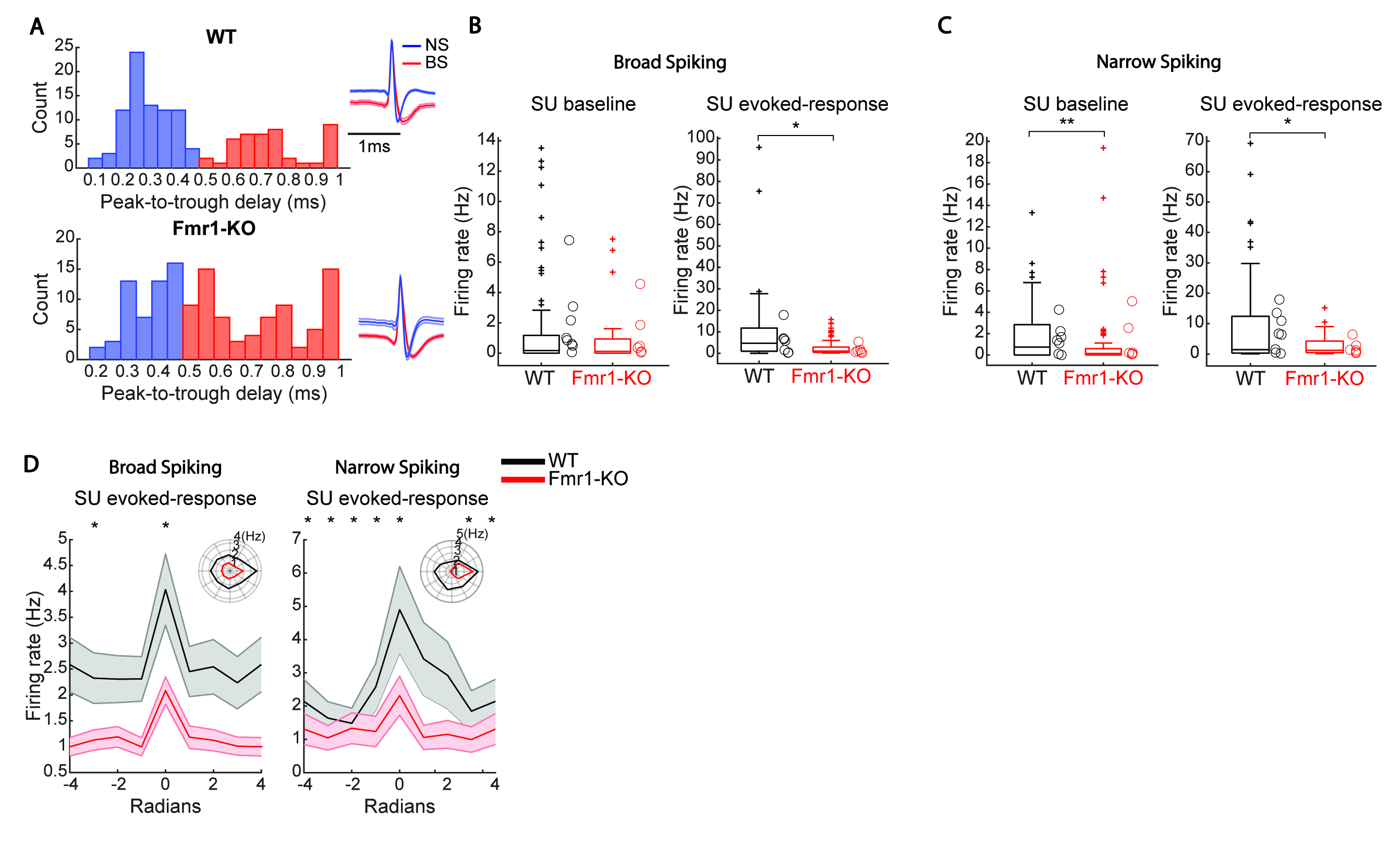

### Supplemental Figure 3

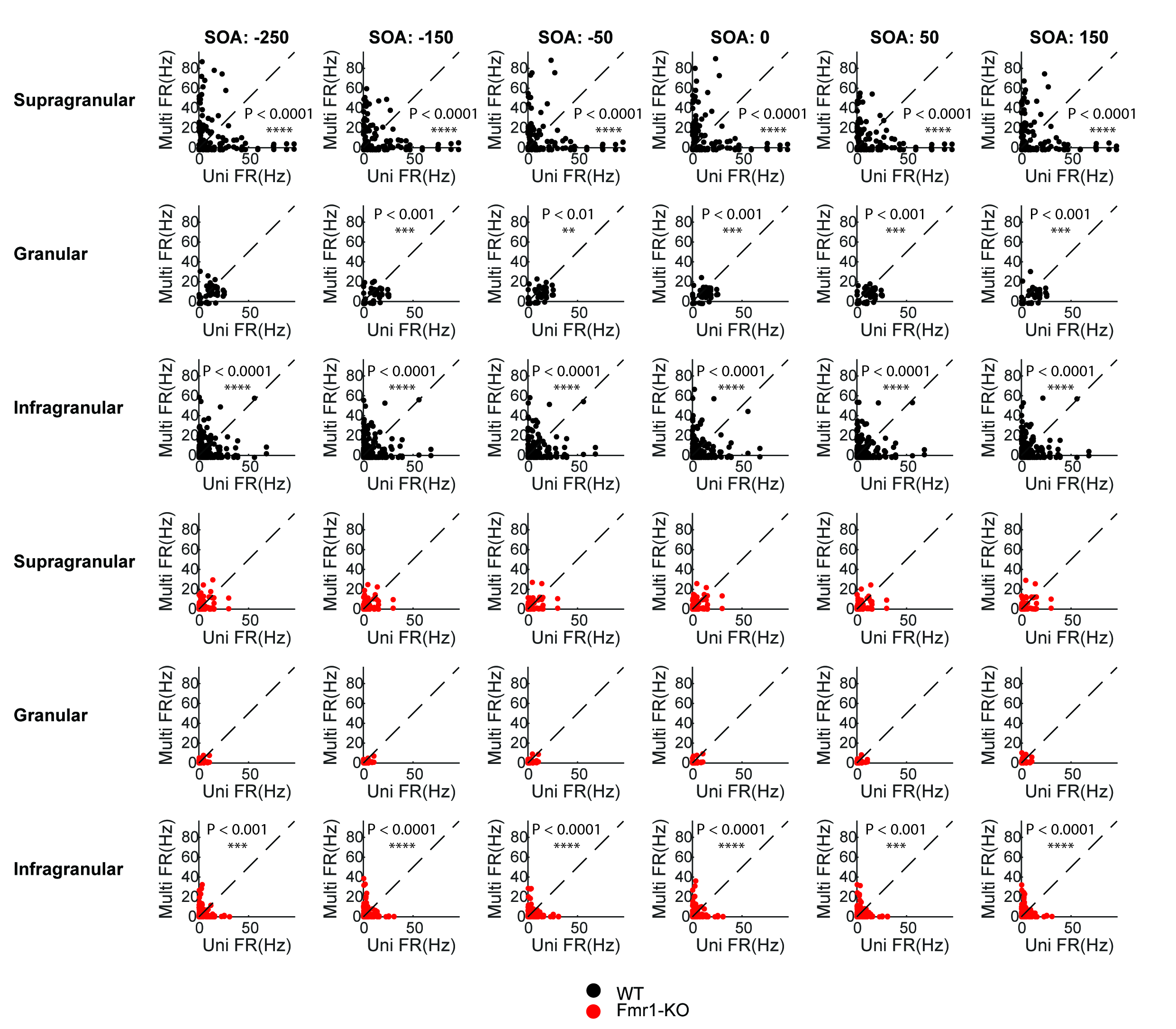

### Supplemental Figure 4

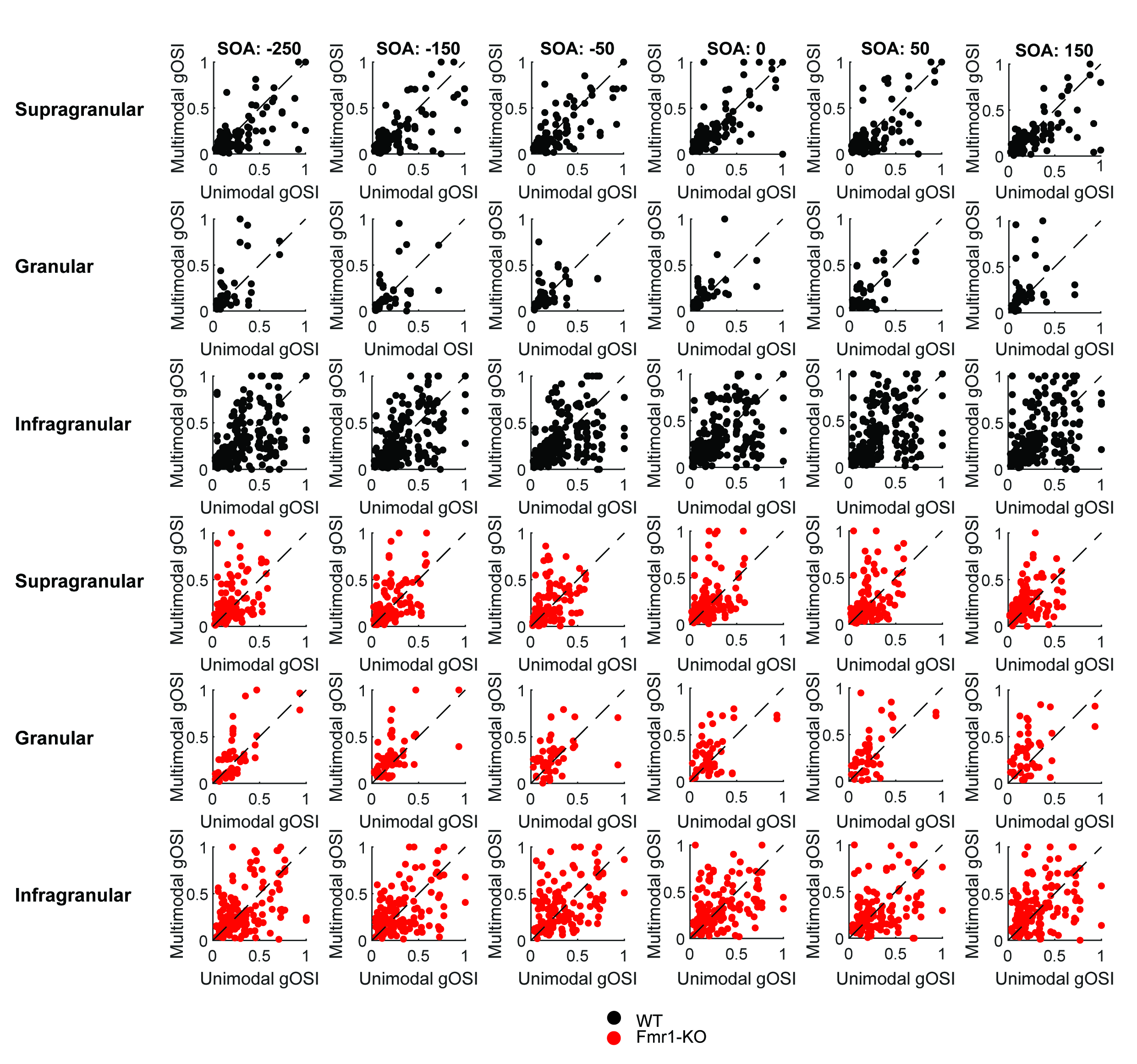

### Supplemental Figure 5

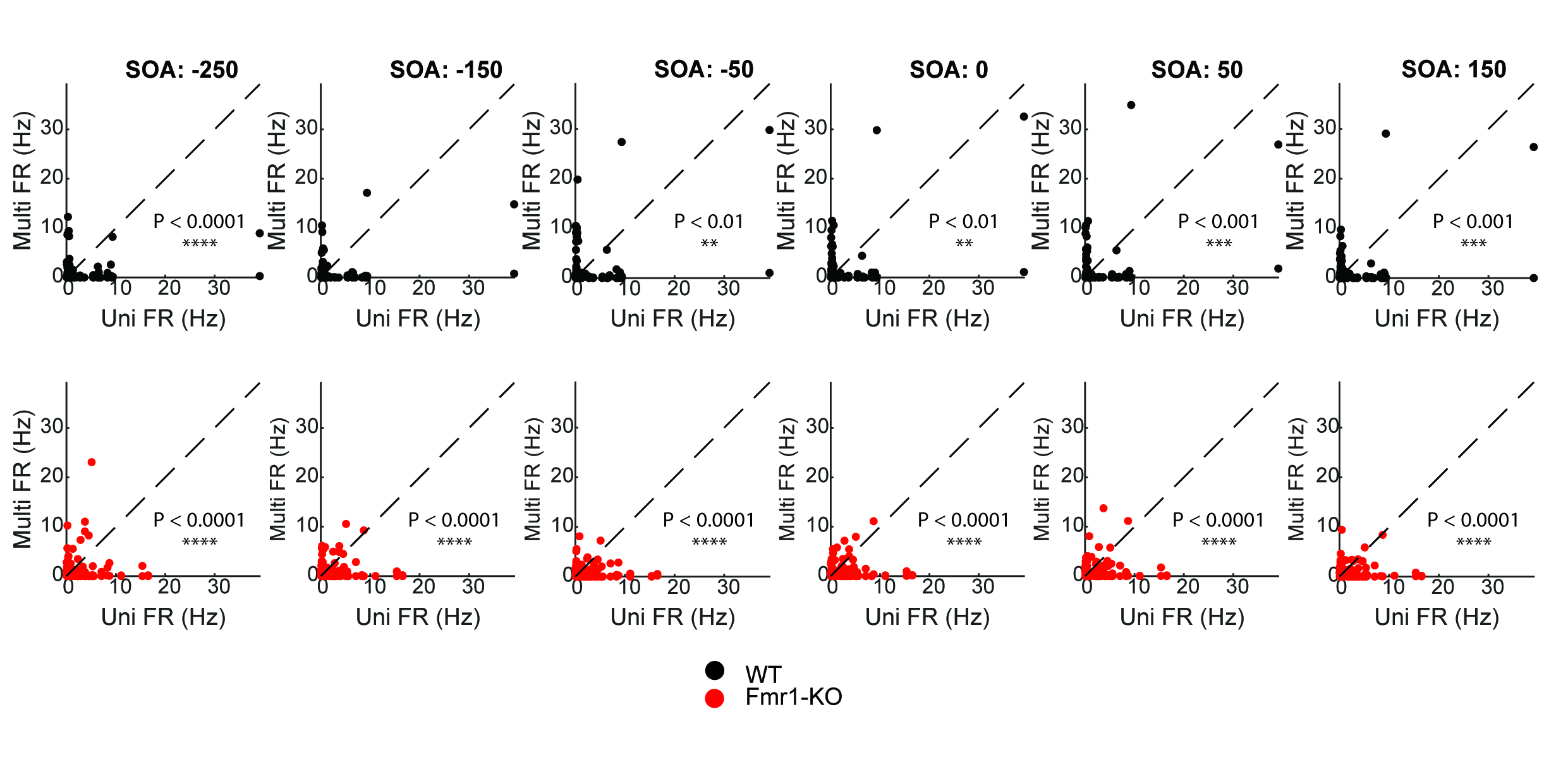
